# Lipid Droplets Direct Amyloid Assembly and β-Cell Stress through Interfacial Control of Amylin Aggregation

**DOI:** 10.64898/2026.06.16.732533

**Authors:** Akash Kumar Jha, Md Asrafuddoza Hazari, Sneha Singh, Gautam Kannan, Shamik Sen, Ashutosh Kumar

## Abstract

Islet amyloid polypeptide (IAPP, amylin) aggregation is a central pathological feature of type 2 diabetes, yet the cellular factors governing its conformational conversion remain incompletely understood. Here, we identify lipid droplets (LDs) as active modulators of amylin structure, aggregation, and β-cell stress. Using artificial and native LDs, we show that amylin binds LD surfaces with high affinity and undergoes accelerated conversion into β-sheet–rich conformations. LDs promote rapid nucleation while constraining fibril elongation, yielding shorter and morphologically distinct amyloid assemblies. Residue-resolved NMR mapping reveals a conserved N-terminal interaction interface, which is amplified upon removal of LD surface proteins, indicating that the LD proteome modulates peptide engagement and aggregation pathways. In β-cells, lipid loading drives intracellular colocalization of amylin with LDs and reshapes transcriptional stress responses, attenuating ER stress and apoptosis while altering markers of β-cell identity. Finally, systemic lipidomic profiling reveals coordinated remodeling of neutral lipid species across dysglycaemic states, linking intracellular LD dynamics with whole-body lipid metabolism. Together, our findings establish lipid droplets as dynamic scaffolds that reshape amylin aggregation pathways and associated β-cell stress responses, providing a mechanistic bridge between lipid dysregulation and islet amyloidosis in diabetes.

**Significance Statement:** Lipid droplets accumulate in pancreatic β-cells during metabolic stress, yet their role in amylin aggregation remains unclear. Using structural, biophysical, and cellular approaches, we show that lipid droplet interfaces directly bind human amylin, reshape its aggregation pathway, and alter fibril morphology. Native lipid droplets and their associated surface proteins further modulate aggregation kinetics and peptide conformations. In β-cells, lipid loading enhances amylin colocalization with lipid droplets and modifies stress and survival responses. Complementary lipidomic profiling reveals systemic remodeling of neutral lipid species across dysglycaemic states. These findings identify lipid droplets as active regulators of amyloid-associated proteostasis, linking lipid dysregulation to β-cell dysfunction in type 2 diabetes.

## Introduction

The pancreatic β-cell plays a central role in maintaining systemic glucose homeostasis through the tightly coordinated co-secretion of insulin and amylin, typically produced in an approximate molar ratio of 100:1 under physiological conditions ^1^. Amylin, or human islet amyloid polypeptide (hIAPP), is a 37-amino-acid peptide of the calcitonin family that contributes to post-prandial satiety, gastric emptying, and glucose regulation via the calcitonin-receptor/RAMP signaling axis ^2,3^. Although amylin is essential for glucose homeostasis, the human variant is intrinsically amyloidogenic, predisposing it to rapid misfolding, oligomerization, and fibril formation under conditions of metabolic stress ^4,5,6^.

Under normal circumstances, β-cell proteostasis is maintained by stringent quality control mechanisms, including the endoplasmic reticulum (ER) stress response pathways, the ubiquitin–proteasome system, and autophagy-lysosomal degradation ^7,8^. These systems collectively act to restrict the buildup of misfolded amylin and prevent intracellular toxicity. However, during insulin resistance and chronic hyperglycemia, which are hallmarks of prediabetes and early type 2 diabetes mellitus (T2DM), β-cells compensate by augmenting insulin secretion. Because insulin and amylin are co-secreted, this adaptive response simultaneously increases amylin biosynthesis. The elevated peptide load, coupled with heightened ER stress, accelerates amylin aggregation kinetics, overwhelms proteostatic pathways, and results in the deposition of toxic oligomers and early fibrillar intermediates ^9,10^.

These early aggregates of hIAPP are known to associate with and permeabilize phospholipid membranes, including the plasma membrane and intracellular organellar membranes, thereby triggering Ca²⁺ dysregulation, ROS generation, mitochondrial stress, and progressive β-cell dysfunction ^11,12^. Such membrane-disruptive species are widely recognized as key drivers of β-cell fragility and apoptosis in type 2 diabetes mellitus (T2DM).

Beyond classical membrane surfaces, emerging evidence highlights a potentially critical yet underexplored role for lipid droplets (LDs) in β-cell pathology. LDs are metabolically active organelles composed of a neutral-lipid core encased in a phospholipid monolayer enriched in perilipin-family proteins and lipid-regulating enzymes ^13^. Recent work suggests that LDs can interact with aggregation-prone proteins, functioning either as protective “sinks” that sequester misfolded species or as catalytic platforms that accelerate amyloid nucleation under lipotoxic conditions ^14^. Their abundance increases markedly in metabolic stress states, such as obesity, lipotoxicity, and type 2 diabetes mellitus (T2DM), with human and rodent studies consistently reporting elevated LD number and enlargement in dysfunctional β-cells ^15–17^. While traditionally regarded as inert lipid reservoirs, LDs are now recognized as dynamic regulators of lipid metabolism, ER–lipid communication, and proteostasis ^18,19^.

This raises an intriguing possibility: given the phospholipid monolayer architecture of LDs and the inherent membrane affinity of toxic hIAPP intermediates, LDs may serve as alternative surfaces that modulate amylin aggregation. Such interactions could, in principle, (i) catalyze aggregation by providing a two-dimensional scaffold, (ii) perturb LD homeostasis and promote aberrant lipid storage, or (iii) disrupt ER–LD communication, thereby altering lipid handling and stress-signaling pathways. Moreover, dysregulated LD biogenesis is linked to altered lipid composition and β-cell dysfunction, suggesting that hIAPP–LD interactions may participate in a self-reinforcing pathological loop that contributes to the progression of T2DM.

Despite these compelling observations, the molecular determinants, kinetics, and functional consequences of hIAPP–LD interplay remain largely unknown. Here, we integrate biochemical, biophysical, and cell-biological analyses to dissect how LDs modulate amylin aggregation and subcellular organization within β-cells. Because systemic lipid availability governs intracellular lipid storage and LD biogenesis, we complemented our cellular analyses with lipidomic profiling across glycaemic states to provide a physiological context for LD-mediated modulation of amylin aggregation. Our findings reveal previously unrecognized modes of crosstalk between lipid storage organelles and amyloidogenic peptides, illuminating a mechanistic axis that may contribute to β-cell vulnerability in metabolic disease.

## Results

### hIAPP binds to artificial lipid droplets and modulates the amyloid assembly kinetics

To establish a mechanistically tractable framework for dissecting hIAPP–lipid droplet interactions, we first employed an artificial lipid droplet (ALD) model system that allows precise control over lipid composition and surface properties. This reductionist approach minimizes the complexity inherent to biological environments and enables isolation of the fundamental physicochemical interactions governing peptide–lipid association, thereby providing a robust basis for extrapolation to physiologically relevant contexts.

After purification of pure monomeric hIAPP and characterization of ALD (Fig. S1A-D), we performed 2D ^15^N–^1^H HSQC nuclear magnetic resonance (NMR) titration experiments. Residue-specific interactions between LMW human islet amyloid polypeptide (hIAPP) and ALDs were probed using chemical shift perturbation (CSP) of 2D resonances. Amylin was titrated with increasing concentrations of ALD over a molar ratio range of 0.1-10 (Fig. 1A, 1B & Fig. S1E), corresponding to pathophysiologically relevant lipid concentrations (30–3000 μM). In the absence of ALD, hIAPP exhibited negligible CSPs, confirming the stability of the reference spectrum. Upon ALD addition, progressive and concentration-dependent CSPs (Fig. 1C-1D) were observed for a distinct subset of residues, while the majority of resonances remained weakly perturbed or unchanged, indicating a selective interaction rather than global unfolding or nonspecific binding. The largest CSPs and intensity reductions were localized to the N-terminal region of hIAPP. Residues R11 and L12 displayed the strongest perturbations, increasing monotonically with ALD concentration. Additional moderate CSPs were detected at residues C2, A5, C7, Q10, A13, and N14, indicating that the N-terminus is the primary ALD interaction interface. In contrast, residues within the central hydrophobic segment, including L16, V17, I26, and L27, exhibited only small but reproducible CSPs that became evident at higher ALD concentrations, consistent with secondary or transient contacts. Several residues, including F15, S19–S20, G24, S28, N31, N35 and T36, showed negligible perturbations under all conditions. Weak CSPs in the C-terminal region, notably at S29 and Y37, and to a lesser extent G33 and S34, were detectable only at high ALD concentrations, suggesting late-stage or low-affinity engagement. Collectively, the CSP profile reveals a highly localized, residue-specific interaction, dominated by the N-terminal segment of hIAPP.

**Figure 1.**
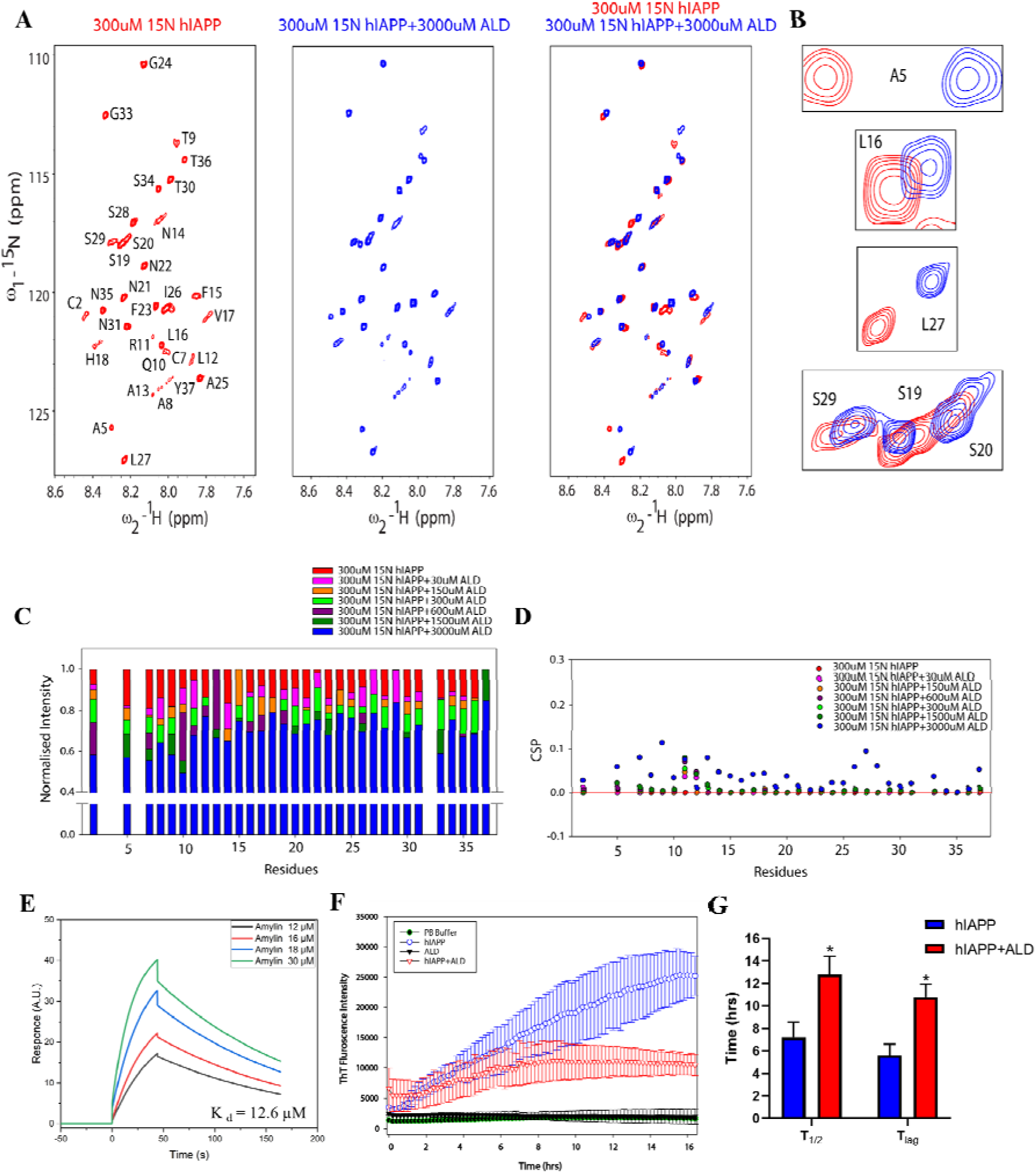
**(A)** ^15^N-^1^H HSQC spectrum illustrating the interaction between LMW human IAPP (hIAPP) in Red with ALD(Blue) in the ratio of 1:10. (**B)** Contour plot of interacting Residues. **(C)** Bar and **(D)** dot plot of Normalized Intensity and CSP of titration of ALD with hIAPP, respectively. **(E)** Surface plasmon resonance (SPR) spectra of hIAPP showing its strong affinity with PMLD. **(F)** ThT Aggregation Kinetics of hIAPP and on interaction with ALD. **(G)** Bar plots show the lag time (T_lag_) and half-time (T_1/2_) of fibril formation upon differential interaction with ALD.

To quantify the affinity of this interaction, surface plasmon resonance (SPR) measurements were performed using immobilized ALDs. Amylin bound ALDs with an equilibrium dissociation constant (K_d_) of 12.6 ± 0.8 μM, indicating a high-affinity interaction consistent with the NMR-derived residue-specific binding pattern (Fig. 1E).

We next assessed the functional consequence of ALD binding on amyloid formation using thioflavin T (ThT) fluorescence kinetics. In the absence of ALD, hIAPP exhibited a half-time of aggregation (T_1/2_) of 7.2 ± 1.4 h and a lag phase (T_lag_) of 5.6 ± 1.01 h. In the presence of ALD, both parameters were significantly prolonged, with T_1/2_ increasing to 13.8 ± 1.6 h and T_lag_ to 10.8 ± 1.12 h. (Fig. 1F & 1G) The comparable standard deviations across conditions indicate that ALDs reproducibly modulate hIAPP aggregation kinetics rather than introducing stochastic effects.

These findings support the possibility that lipid droplet interfaces can modulate hIAPP assembly, as ALDs bind hIAPP via a selective N-terminal interface, form a high-affinity complex, and significantly reshape the temporal progression of amyloid formation.

### Lipid droplet model system attenuates **β**-sheet conversion of hIAPP and generates structurally distinct amyloid fibrils

To explore how minimal reconstituted lipid droplet model systems influence the structural evolution of hIAPP during amyloidogenesis, we characterized secondary structure transitions and aggregate morphology in the presence and absence of ALDs using spectroscopic and nanoscale imaging approaches. In solution, hIAPP underwent a gradual transition from predominantly disordered or α-helical conformations to β-sheet–rich structures, consistent with the canonical amyloidogenic pathway. In contrast, co-incubation with ALDs markedly altered this trajectory. Circular dichroism spectra revealed a pronounced reduction in β-sheet content accompanied by a substantial increase in α-helical signal (Fig. 2B & 2C). These changes suggest that ALD surfaces stabilize aggregation-prone conformations by providing hydrophobic interfaces that facilitate peptide alignment and β-strand stacking, a mechanism similar to those reported for other lipid-mediated amyloid systems. Compared with hIAPP in solution, the zwitterionic DOPC: DOPE monolayer on synthetic lipid droplets significantly delays the α-helix-to-β-sheet transition and subsequent aggregation of hIAPP. While membrane surfaces generally enhance amyloid formation by functioning as nucleation platforms, our results reveal that lipid surface chemistry is a significant factor in aggregation kinetics.

**Figure 2.**
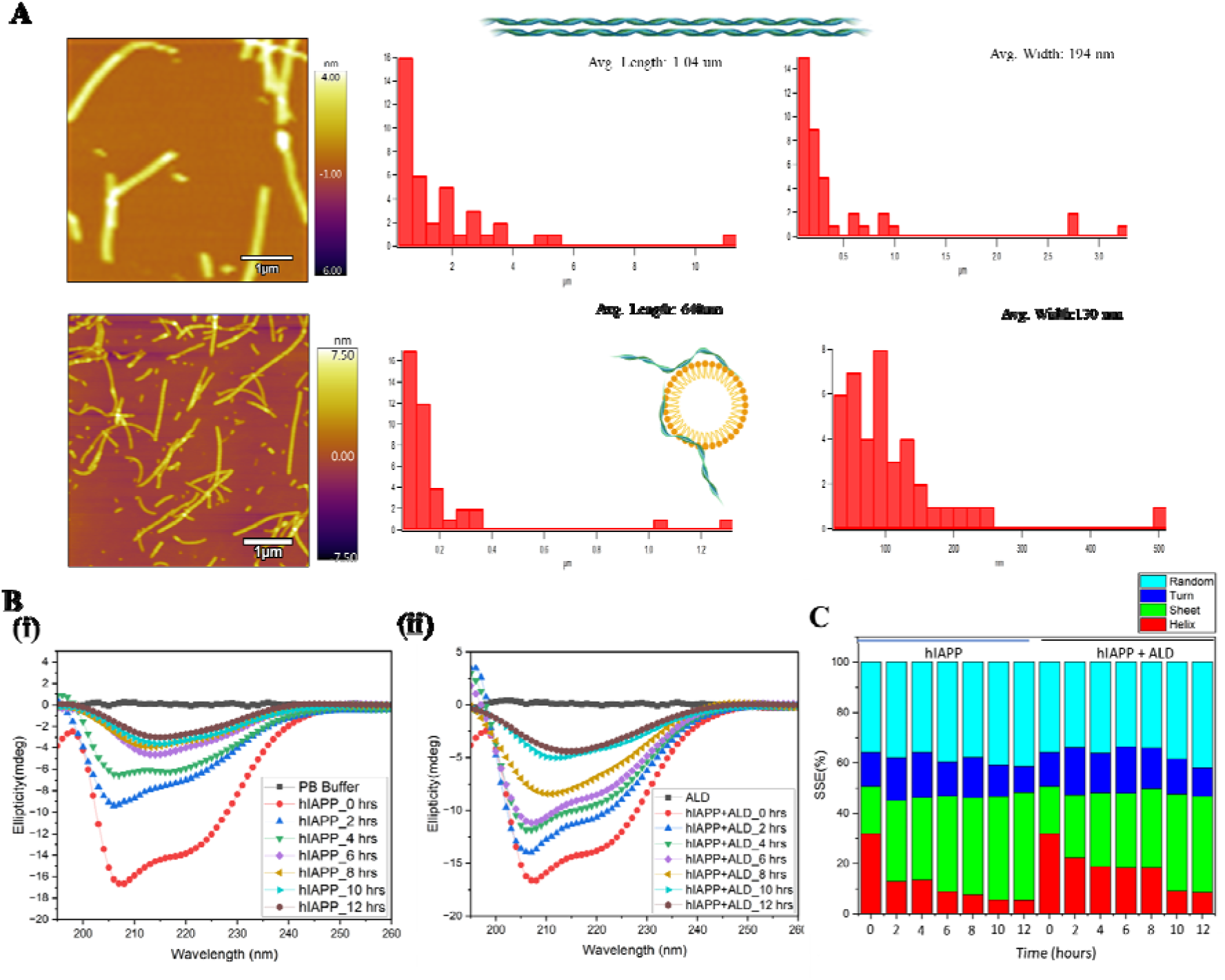
**(A)** Atomic force microscopy (AFM) images and fibril dimension analysis of hIAPP aggregates. Representative AFM images show fibrils formed in the absence (upper panel) and presence (lower panel) of ALDs. **(B)** & **(C)** Circular dichroism (CD) analysis of secondary structural transitions in hIAPP in the absence and presence of artificial lipid droplets (ALD): Time-dependent changes in α-helix, β-sheet, turn, and random coil content were quantified from CD spectra of hIAPP alone and hIAPP incubated with ALD over a 12-hour period.

Next, we visualized endpoint fibrils using atomic force microscopy to assess whether these conformational effects resulted in a particular morphology of aggregate architecture or not (Fig. 2A). In the absence of ALDs, hIAPP formed long, mature fibrils with an average length of approximately 1.04 µm and a mean width of ∼192 nm, consistent with typical amyloid morphology. By contrast, fibrils formed in the presence of ALDs were markedly shorter (∼640 nm) and thinner (∼130 nm), indicating a substantial shift in the aggregation pathway. The pronounced reduction in fibril length suggests that ALDs function as heterogeneous nucleation platforms that accelerate early aggregation while constraining subsequent elongation. This may arise from sequestration of growing fibril ends at the droplet interface, steric hindrance imposed by droplet curvature, or altered monomer–fibril association dynamics. The decreased fibril width further implies impaired lateral association of protofibrils or the formation of structurally distinct polymorphs.

Together, these data demonstrate that within a reductionist LD-interface model, the conformational conversion of hIAPP toward β-sheet–rich assemblies also fundamentally remodels fibril architecture. By biasing aggregation toward shorter, thinner fibrils, lipid droplets reshape the structural and morphological landscape of amylin amyloids, with potential implications for their biological activity and cytotoxicity.

To determine whether the interaction principles observed in ALDs extend to biologically derived droplets, we next examined hIAPP engagement with native rat liver LDs.

### LD surface proteins modulate hIAPP binding and aggregation dynamics

Building on our observation that artificial lipid droplets reshape the conformational and morphological landscape of hIAPP aggregates, we next investigated whether biologically derived lipid droplets (LDs) and their associated surface proteins similarly regulate amylin interactions. To delineate the specific contribution of LD-bound proteins, we compared hIAPP binding to intact LDs with that to proteinase K-treated LDs (pkLDs), in which surface proteins were selectively removed while preserving droplet integrity.

Droplet concentration was normalized across artificial lipid droplets (ALDs), LDs, and pkLDs to exclude differences in droplet abundance as a confounding variable (Fig. S2A). Proteinase K treatment efficiently depleted LD-associated surface proteins, yielding pkLDs suitable for interaction studies. Dynamic light scattering and cryo-TEM analyses confirmed that protease treatment did not compromise droplet morphology or induce aggregation (Fig. S2B & S2C). The resulting particles displayed a mean hydrodynamic radius of 185 ± 55 nm and a modest polydispersity index of 0.6 ± 0.2, indicating a relatively homogeneous population appropriate for biophysical characterization.

Residue-specific interactions were mapped by 2D ^15^N–^1^H HSQC chemical shift perturbation (CSP) analysis upon titration of hIAPP with increasing concentrations of LDs or pkLDs. In the absence of lipid assemblies, hIAPP exhibited minimal CSPs, confirming spectral stability. Addition of intact LDs induced modest but concentration-dependent CSPs, predominantly localized to the N-terminal region (residues 2–14) (Fig.3A–3C). Residues C2, A8, R11, L12, and A13 showed progressive increases in CSP magnitude, reaching maximal perturbation at 1500–3000 µM LD. Smaller and delayed perturbations were also observed in central residues (L16, V17, H18) and in the C-terminal residue Y37. In contrast, titration with pkLDs produced substantially larger CSP amplitudes across the same residue set. At higher pkLD concentrations, pronounced perturbations were detected for residues C2, C7, A8, R11, L12, A13, H18, and Y37, with several residues exhibiting a sharp increase beyond 600 µM pkLD. Although the magnitude and dynamic range of CSPs were consistently greater for pkLDs than for intact LDs, the overall spatial distribution of affected residues remained conserved, indicating a common interaction interface. Consistent trends were observed in complementary intensity analyses, where pkLDs caused stronger signal attenuation or modulation, particularly at elevated concentrations.

**Figure 3.**
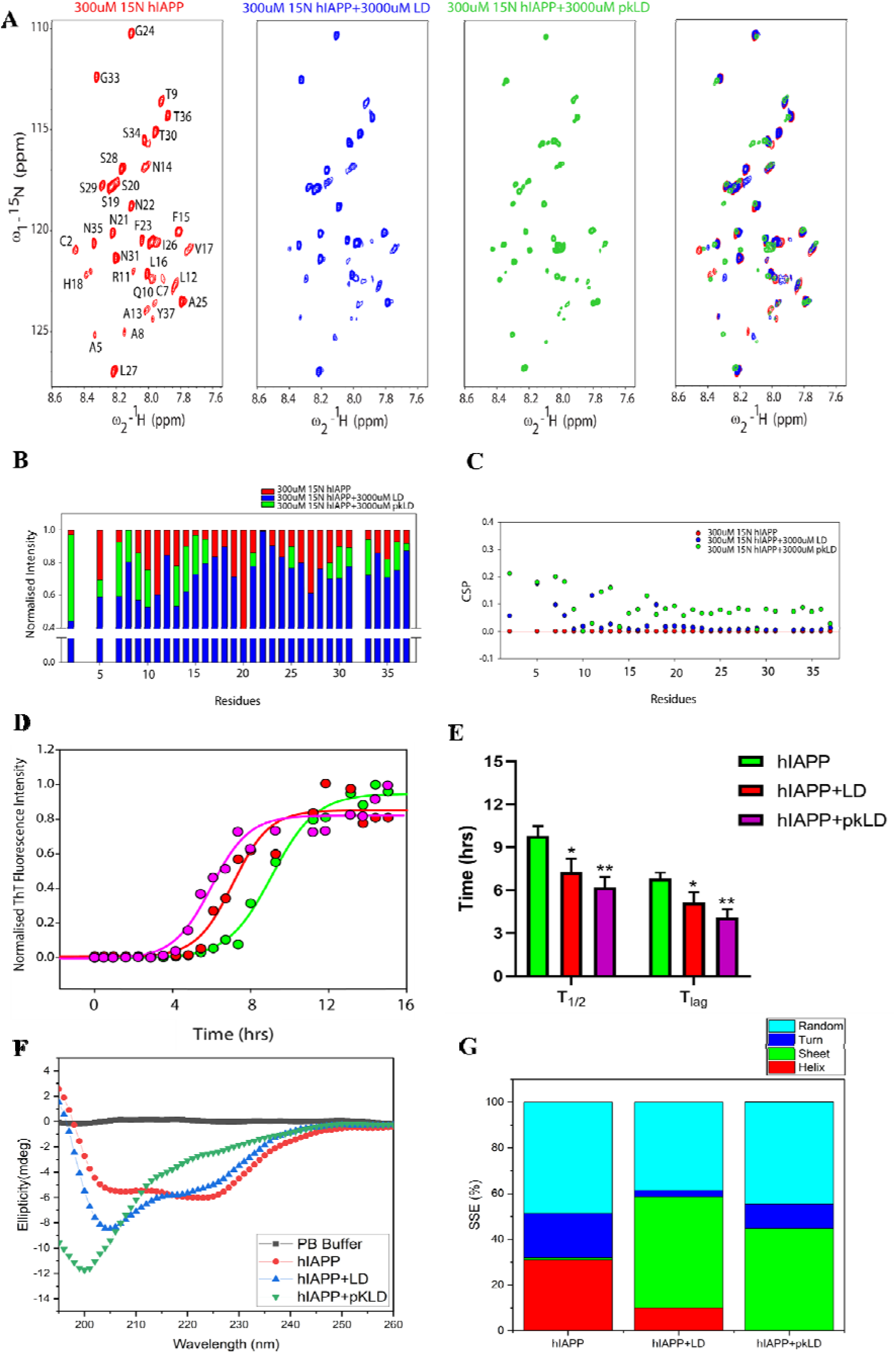
**(A)** ^15^N HSQC spectrum illustrating the interaction between human IAPP (hIAPP) in Red with LD (Blue) and pkLD (light green) in the ratio of 1:10. (**B)** Bar and (**C)** dot plot of Normalized Intensity and CSP, respectively, in the lower panel showing comparative interaction of hIAPP with LD and pkLD. (**D)** ThT Aggregation Kinetics of hIAPP and on interaction with LD and pkLD. (**E)** Bar plots show the lag time (T _lag_) and half-time (T_1/2_) of fibril formation upon differential interaction with LD and pkLD. **(F)** CD spectra of hIAPP and hIAPP titrated sample after HSQC-NMR recording, and **(G)** their respective secondary structure estimation.

We next assessed the functional consequences of these interactions on amyloid formation using Thioflavin T (ThT) aggregation kinetics. In the absence of lipid droplets, hIAPP aggregated with a half-time (T₁/₂) of 9.8 ± 0.8 h and a lag time (T _lag_) of 6.84 ± 0.4 h. The presence of intact LDs markedly accelerated aggregation, reducing T₁/₂ to 6.4 ± 1.2 h and T _lag_ to 4.12 ± 0.7 h. Protease-treated LDs also promoted aggregation relative to hIAPP alone (T₁/₂ = 7.2 ± 1.1 h; T _lag_ = 5.76 ± 1.2 h), but to a lesser extent than intact LDs. The kinetic growth parameter (b) followed a similar trend, being highest for hIAPP alone (1.48), intermediate for hIAPP + LD (1.14), and lowest for hIAPP + pkLD (0.72), indicating altered fibril growth dynamics in the presence of lipid droplets and their surface proteins (Fig. 3D & 3E).

Far-UV circular dichroism spectroscopy further revealed distinct conformational effects of LDs and pkLDs on hIAPP (Fig. 3F & 3H). Free hIAPP adopted a largely disordered ensemble, with high random coil content (∼49%), moderate α-helicity (∼31%), and minimal β-sheet structure. Interaction with intact LDs induced a pronounced conformational shift toward amyloidogenic states, characterized by a substantial increase in β-sheet content (∼49%) accompanied by a reduction in α-helical and random coil contributions. In contrast, pkLDs abolished α-helicity, increased turn and random coil content, and promoted β-sheet formation to a lesser extent (∼45%), consistent with a more heterogeneous and less ordered structural ensemble.

Together, these findings demonstrate that, while both intact and protein-depleted lipid droplets engage hIAPP via a conserved interaction interface, LD-associated surface proteins significantly modulate binding strength, aggregation kinetics, and conformational outcomes. Intact LDs preferentially stabilize β-sheet–rich amyloidogenic conformations and accelerate fibril formation, whereas pkLDs bias hIAPP toward more disordered, intermediate states. This indicates that LD surface proteins play an active role in shaping the structural and kinetic landscape of amylin aggregation.

### Lipid loading promotes intracellular amylin colocalization

To investigate the interaction of amylin with lipid droplets (LDs) and its impact on β-cell viability, we used INS-1E cells with tetracycline-inducible GFP-tagged amylin expression and pharmacological induction of LD formation. Analysis suggested that combined treatment with tetracycline (TA) and oleic acid (OA) led to a marked increase in cell death compared with either treatment alone (Fig. S3A-S3C). Whereas TA or OA individually caused only minimal cytotoxicity, their combination resulted in a marked rise in propidium iodide (PI)–positive cells, indicating synergistic loss of membrane integrity and reduced viability. These findings suggest that lipid loading sensitizes β-cells to amylin-induced stress, or conversely, that amylin expression exacerbates lipid-mediated toxicity. While the live–dead assay demonstrates enhanced cytotoxicity under combined lipid-loading and amylin-expression conditions, the present study does not resolve which specific aggregate species mediate this effect.

Confocal imaging of INS-1E cells stained with LipidTOX dye demonstrated extensive intracellular LD formation following OA treatment and revealed prominent spatial overlap between LDs and GFP-tagged amylin (Fig. 4). Quantitative colocalization analysis using Pearson’s correlation coefficient yielded a value of 0.665, consistent with substantial colocalization of amylin with LDs (Fig. 4D). Notably, detectable colocalization was also observed in OA-treated cells lacking TA induction, indicating preferential association of basal amylin with lipid-rich subcellular compartments. These data support the existence of a physical interaction between intracellular LDs and amylin in living β-cells.

**Figure 4.**
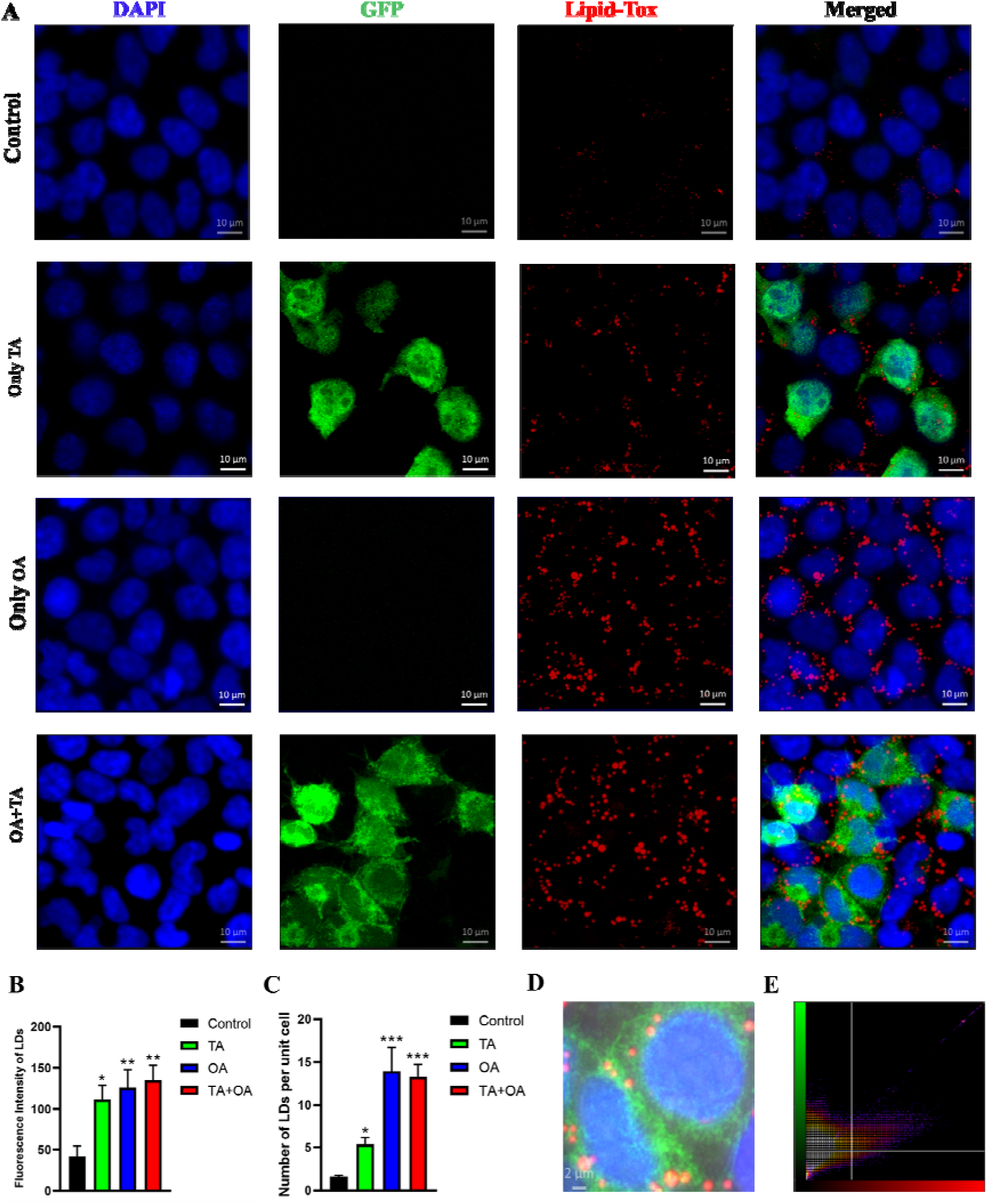
**(A)** Confocal fluorescence microscopy showing GFP expression and lipid droplet accumulation in INS-1 amylin overexpressing cells. Columns show nuclear staining with DAPI (blue), intrinsically GFP-labeled hIAPP (green), Lipid-Tox staining of neutral lipid droplets (red), and merged images, respectively. Oleic acid (OA) enhanced LD Production, and Tetracycline (TA) enhanced Amylin production. Scale bar: 10 µm. **(B)** Change in LD intensity across different treatment conditions. **(C)** Number of LDs per unit cell at different treatment conditions. *PO<O0.05, **PO<O0.01, ***PO<O0.001. **(D)** Enlarged image (2µm) of TA+OA-treated cells showing Colocalization with a Pearson correlation coefficient of 0.665

Quantitative analysis of LD fluorescence intensity showed a strong treatment-dependent increase in neutral lipid accumulation. Control cells displayed low LD intensity (41.87□±□12.96), reflecting basal lipid storage. Tetracycline (TA) treatment significantly increased LD intensity to 111.68□±□17.26, while OA treatment further elevated LD intensity to 125.90□±□21.77. Combined TA and OA treatment produced the highest LD intensity (135.00□±□18.42), suggesting additive or synergistic enhancement of LD biogenesis (Fig. 4B).

Consistent with these findings, LD number per cell was markedly increased under lipid-loading conditions. Control cells exhibited a low basal LD count (1.60□±□0.16 droplets per cell). TA treatment induced a greater than threefold increase (5.37□±□0.82), whereas OA produced the most pronounced effect (13.94□±□4.75). Combined TA and OA treatment also significantly elevated LD abundance (13.25□±□1.42), although to a slightly lower extent than OA alone, indicating potential competition between lipid species for storage capacity (Fig. 4C).

### Coupled lipid storage and IAPP expression reprogram **β**-cell stress pathways

To assess the cellular consequences of lipid droplet (LD)–amylin interactions, we examined transcriptional responses in β-cells subjected to inducible amylin expression in the presence or absence of oleic acid (OA), which promotes LD biogenesis. Gene expression profiling by RT–qPCR revealed a coordinated stress response to amylin overexpression, characterized by activation of endoplasmic reticulum (ER) stress markers, apoptotic pathways, and alterations in β-cell identity factors.

Amylin induction alone led to a pronounced upregulation of unfolded protein response (UPR) components and apoptotic mediators, consistent with proteotoxic stress. Transcripts encoding ER stress regulators and executioner caspases were significantly elevated, indicating engagement of both adaptive and pro-apoptotic branches of the UPR. In parallel, expression of key β-cell transcription factors was altered, reflecting disruption of β-cell functional identity under conditions of amyloidogenic stress (Fig. 5B).

**Figure 5.**
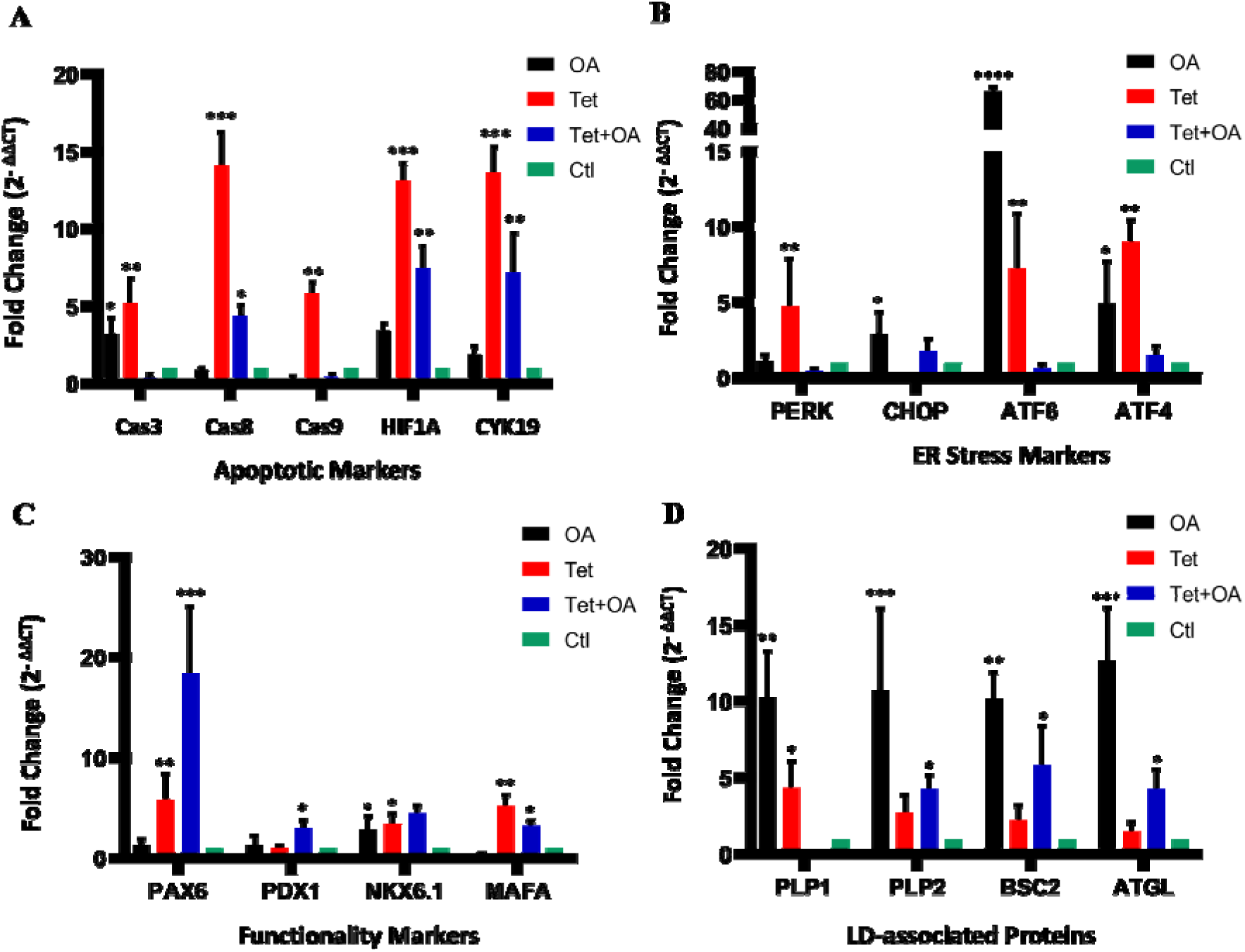
RT-PCR analysis of **(A)** Apoptotic and stress-responsive genes. **(B)** ER stress markers. **(C)** Functionality markers, and **(D)** Lipid Droplet-associated protein markers in INS-1E at different treatment conditions. INS-1E cells were treated with tetracycline (Tet), oleic acid (OA), or both (Tet + OA). Red bars represent Tet-induced amylin overexpression, black bars represent OA treatment to promote lipid droplet accumulation, and blue bars indicate combined Tet + OA treatment. Untreated control cells are shown in green. mRNA expression levels were quantified and normalized to the housekeeping GAPDH gene. Data represent mean ± SEM of triplicates

Notably, co-treatment with OA substantially modified this transcriptional landscape. OA-induced LD formation attenuated the expression of several ER stress and apoptotic markers compared with amylin induction alone, suggesting partial mitigation of the proteotoxic burden. This reduction in stress signaling coincided with a marked increase in expression of LD-associated genes, including *PLIN2* and *BSCL2*, confirming effective induction of the LD biogenesis program. These findings are consistent with a role for LDs as buffering compartments that sequester misfolded or aggregation-prone proteins, thereby limiting their interaction with ER membranes and chaperone systems (Fig. 5D).

Analysis of β-cell identity genes further revealed a complex adaptive response. Amylin overexpression led to upregulation of PAX6, PDX1, and NKX6.1, suggesting the activation of compensatory transcriptional programs aimed at preserving the β-cell phenotype. In contrast, expression of *MAFA* was suppressed under lipid co-treatment, indicative of early β-cell dysfunction or dedifferentiation, a hallmark of progressive type 2 diabetes. These divergent transcriptional changes imply that LD formation reshapes not only stress signaling but also the regulatory networks governing β-cell identity (Fig. 5C).

Additional stress-associated pathways were also modulated. *HIF1A* expression was elevated in both amylin-induced and amylin + OA conditions, consistent with activation of pseudo-hypoxic signaling commonly observed under mitochondrial or ER stress. Conversely, OA treatment suppressed expression of cytoskeletal regulator *CYK19*, suggesting potential stabilization of cytoskeletal architecture and organelle positioning during lipid adaptation (Fig. 5A).

Together, these data demonstrate that LD biogenesis significantly alters the cellular response to amyloidogenic amylin. While amylin overexpression elicits strong ER stress and apoptotic signaling, concomitant LD formation partially alleviates this proteotoxic stress while simultaneously reprogramming β-cell transcriptional identity. These findings indicate that LDs function as active modulators of proteostasis and stress signaling in β-cells, linking lipid metabolism to the cellular handling of amyloidogenic proteins.

**Figure 6.**
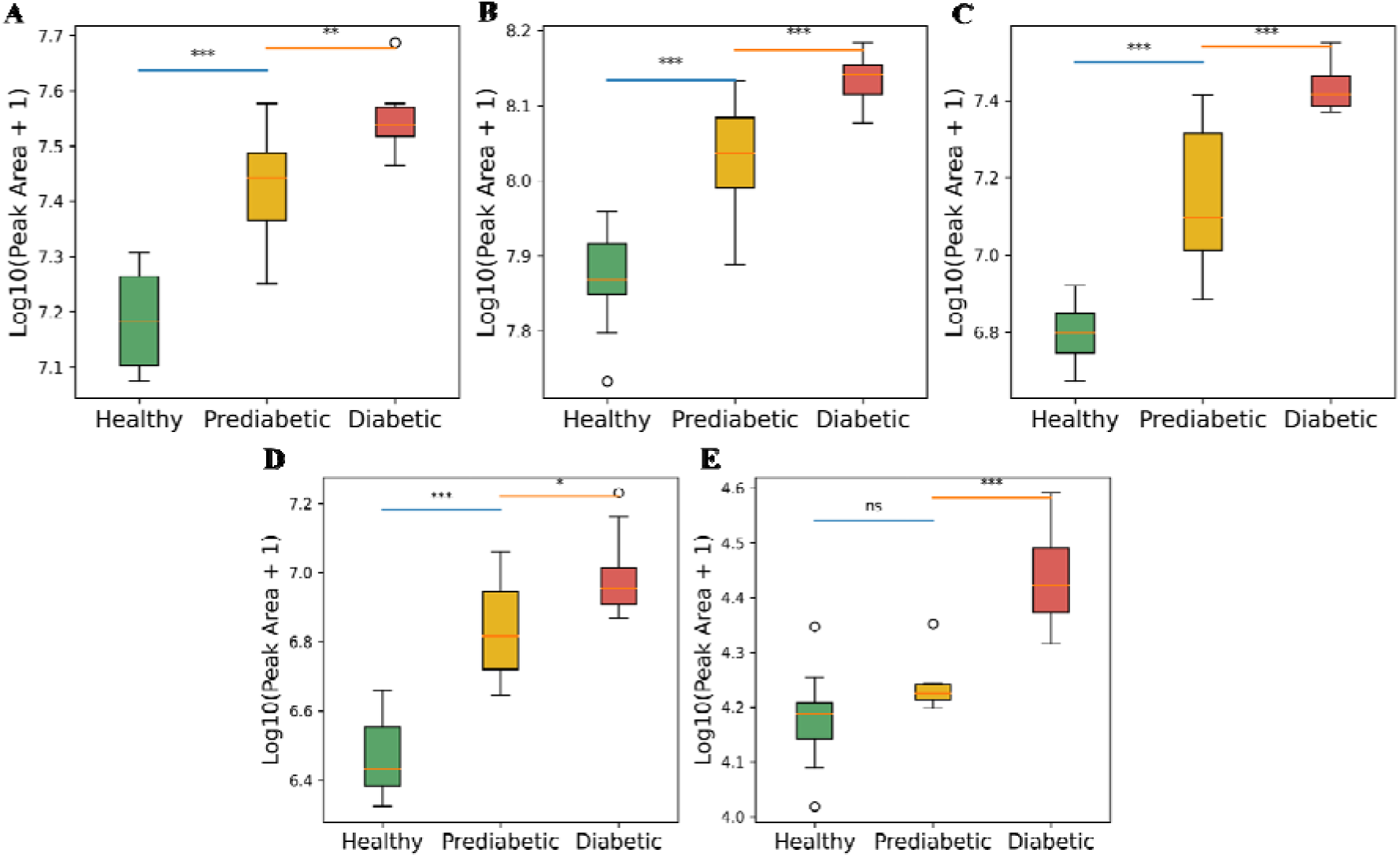
Altered neutral lipid profiles across healthy, prediabetic, and diabetic individuals. Box-and-whisker plots showing log10-transformed peak areas of selected neutral lipid species in healthy, prediabetic, and diabetic cohorts. (A) TG 18:0_18:1_18:2, (B) TG 16:0_18:1_18:2, (C) DG 18:1_18:2, (D) DG 18:2_18:2, and (E) CE 20:0. Boxes represent the interquartile range with the median indicated by the center line; whiskers denote the minimum and maximum values. Statistical significance between groups was assessed using a two-sided Mann-Whitney U test. A progressive increase in triglyceride, diacylglycerol, and cholesteryl ester species is observed from healthy to diabetic states, indicating dysregulated neutral lipid metabolism during the progression of metabolic disease. *PO<O0.05, **PO<O0.01, ***PO<O0.001.

## Discussion

This study identifies lipid droplets as active modulators of amylin aggregation and links cellular lipid–amyloid interactions to systemic remodeling of neutral lipid metabolism under dysglycaemic conditions. By integrating residue-specific binding, aggregation kinetics, conformational analysis, fibril morphology, transcriptional responses, and circulating lipidomics, we establish lipid droplets as interfacial platforms that reshape amyloid pathways and metabolic stress responses.

We acknowledge that the artificial lipid droplets used here do not fully reproduce the compositional and proteomic complexity of native β-cell LDs. Rather than serving as physiological replicas, ALDs were employed as reductionist systems to isolate the contribution of phospholipid monolayer interfaces to hIAPP interaction dynamics. Human islet amyloid polypeptide (hIAPP) engages lipid droplets through a localized and residue-specific interface dominated by its N-terminal region, consistent with amphipathic recognition of lipid–water interfaces and with prior reports identifying this segment as a primary membrane-contact site ^49^. PC/PE mixed membranes create a densely packed, ordered interface with fewer transitory hydrophobic defects by limiting water penetration into the hydrophobic core while improving headgroup hydration. These structural characteristics limit partial peptide insertion, which is necessary for nucleating β-sheets. As a result, hIAPP probably remains highly hydrated and weakly associated at the membrane surface, which is detrimental to the low-polarity environment required for rapid β-sheet stabilization. This interaction stabilizes a population of surface-associated conformers without globally unfolding the peptide. Rather than simply increasing peptide adsorption, lipid droplets selectively redistribute aggregation-competent species, thereby altering the temporal progression of amyloid formation ^50,51^. The prolongation of the lag and growth phases, together with enhanced β-sheet accumulation, indicates that lipid droplets promote conformational conversion while constraining fibril elongation ^52,53^.

Consistent with this interpretation, atomic force microscopy revealed marked remodeling of fibril morphology in the presence of lipid droplets, resulting in shorter, thinner aggregates. Such structural features are characteristic of heterogeneous nucleation on surfaces, where templated assembly accelerates early aggregation steps but limits longitudinal and lateral growth ^54,55^. A further reduction in fibril width suggests impaired protofibril association or the emergence of distinct polymorphic forms. Although LDs altered fibril morphology and aggregation kinetics, the present study does not establish whether the resulting assemblies are more or less membrane-disruptive at the molecular level. Because fibril dimensions can influence surface reactivity and membrane interactions, LD-induced remodeling of hIAPP fibrils may contribute to the altered cytotoxic and stress responses observed in β-cells ^56,57^.

Native lipid droplets further modulate these processes through the surface proteins associated with them. Residue-resolved NMR revealed a conserved interaction interface centered on the N-terminus of hIAPP, yet protease-treated droplets produced substantially larger chemical shift perturbations, indicating enhanced peptide–lipid contact upon removal of surface proteins. Despite this increased binding, intact lipid droplets accelerated aggregation more efficiently and stabilized β-sheet–rich conformations ^58,59^. This divergence indicates that LD surface proteins do not merely shield lipid interfaces but actively template productive amyloidogenic pathways. In contrast, protein-depleted droplets promoted structurally heterogeneous, intermediate conformations with reduced α-helicity, consistent with altered coupling between nucleation and elongation steps ^60,61^. Although the specific LD-associated proteins responsible for modulating hIAPP interaction were not identified in the present study. Future proteomic and reconstitution-based approaches will be required to resolve the molecular determinants governing this effect.

At the cellular level, lipid droplet biogenesis modulates stress responses elicited by amyloidogenic amylin. Induction of amylin alone activates the unfolded protein response and apoptotic pathways, consistent with its established proteotoxicity in β-cells. Promoting lipid droplet formation attenuates several of these stress markers while upregulating genes associated with droplet biogenesis, supporting a buffering role for lipid droplets against proteostatic stress. This is consistent with emerging evidence that lipid droplets can sequester misfolded or aggregation-prone proteins, thereby limiting their interaction with vulnerable membranes and organelles. However, lipid co-treatment also suppresses key β-cell identity factors, indicating that stress mitigation may occur at the expense of functional differentiation and may contribute to progressive β-cell dysfunction ^62,63,64^.

The induction of pseudo-hypoxic signaling in both amylin-expressing and lipid-treated cells further suggests convergence between proteotoxic and metabolic stress pathways. Such coupling likely reflects shared perturbations of mitochondrial and endoplasmic reticulum homeostasis, reinforcing the concept that amyloid burden and lipid metabolism are mechanistically intertwined in β-cell pathology ^65,66^.

Extending these findings to the systemic level, we demonstrate that dysglycaemia is associated with coordinated remodeling of circulating neutral lipid species. Because oleic acid independently remodels ER stress signaling and lipid metabolism, the transcriptional effects observed under OA treatment cannot be exclusively attributed to direct LD–amylin interactions. Instead, the data support a model in which lipid loading modifies the cellular context in which amylin proteotoxicity occurs. Glycemia-associated remodeling of circulating triglyceride (TG), diacylglycerol (DG), and cholesteryl ester (CE) species observed in this study provides a systemic framework for the lipid droplet (LD)–amylin axis depicted in Fig. 7.

**Figure 7.**
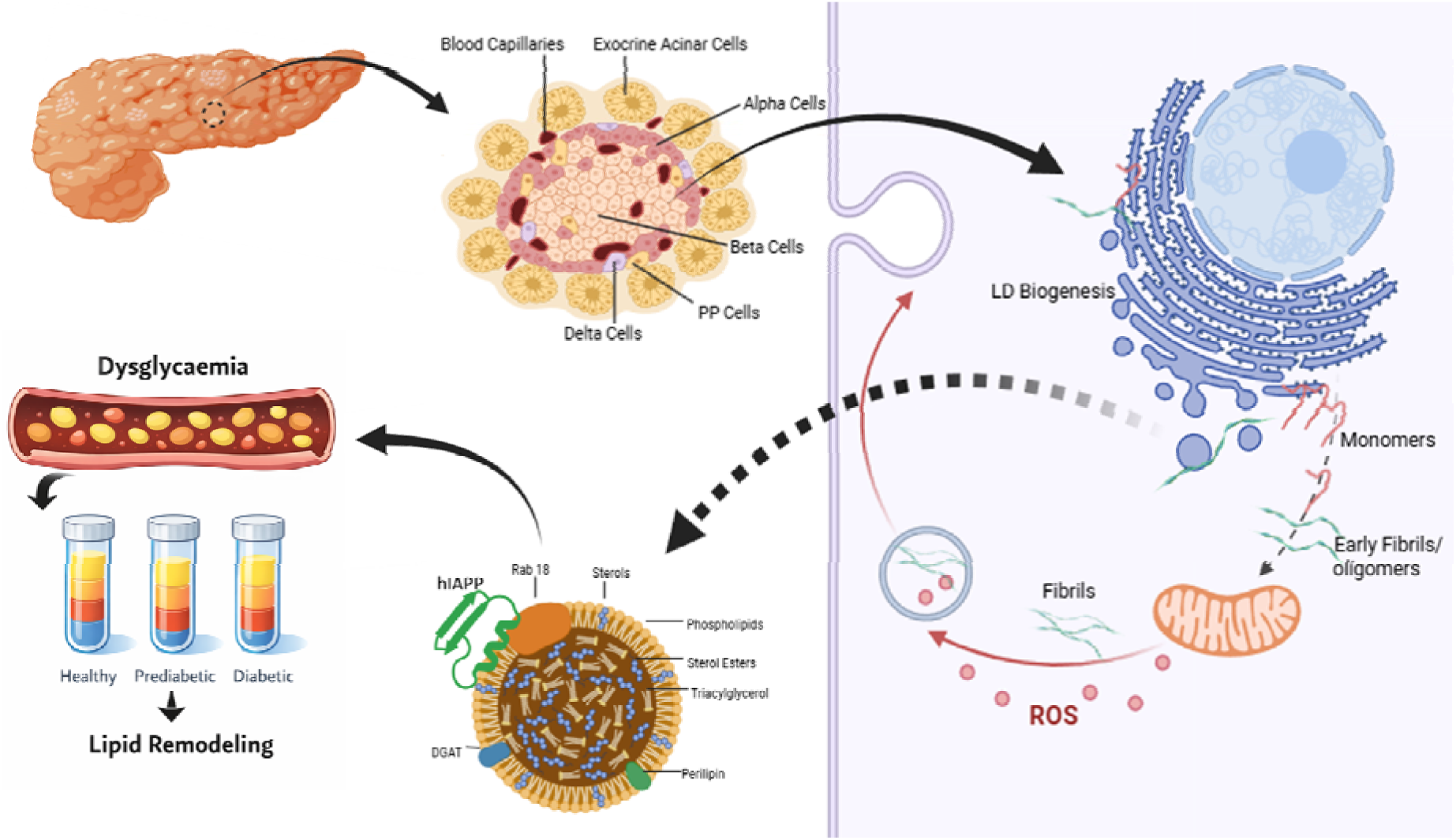
Model illustrating lipid droplet (LD)–dependent modulation of amylin aggregation and associated. β**-cell stress**. Metabolic stress increases LD biogenesis and intracellular amylin (or hIAPP), enabling LD surfaces to act as high-affinity scaffolds that promote amylin β-sheet conversion, rapid nucleation, and formation of short fibrils. LD-associated aggregates induce ER stress, disrupt proteostasis, and reduce the number of mature insulin granules. Concurrently, amylin stress triggers compensatory LD expansion, reflecting altered lipid metabolism aimed at limiting lipotoxicity. However, increased LD abundance provides additional amyloid-reactive interfaces, establishing a maladaptive feedback loop that exacerbates β-cell dysfunction. This LD–amylin–stress axis links local proteotoxicity with impaired insulin secretion and systemic metabolic dysregulation in type 2 diabetes mellitus.

Neutral lipids in circulation are not only biomarkers of metabolic dysfunction but also major determinants of lipid flux into peripheral tissues, where they drive intracellular lipid storage and LD biogenesis^67,68^. Elevated DG species are of particular pathophysiological relevance because they activate protein kinase C signaling pathways that impair insulin action and exacerbate β-cell secretory demand, thereby increasing amylin production^69,70^. In parallel, excess TG- and CE-rich lipoproteins promote ectopic lipid deposition in non-adipose tissues, including pancreatic islets, leading to LD expansion and lipotoxic stress^71^.

Emerging evidence further indicates that LDs function as dynamic organelles involved in proteostasis and can sequester or modulate aggregation-prone proteins^72,73^. Within this context, our cellular data demonstrate that LD abundance and surface properties reshape amylin aggregation pathways and stress responses, suggesting that lipid oversupply may indirectly influence amyloid toxicity by altering intracellular lipid storage capacity. Thus, systemic lipidomic alterations and LD-dependent modulation of amylin proteotoxicity are best interpreted as mechanistically coupled processes operating across biological scales rather than independent consequences of hyperglycemia. Physiologically, this coupling may constitute a maladaptive feedback loop in which lipid oversupply drives LD formation, LDs alter amylin aggregation, and the resulting proteotoxic stress further compromises β-cell function and whole-body metabolic homeostasis.

Overall, our results support a model in which LDs act as high-affinity scaffolds that capture soluble amylin, accelerate its structural conversion to β-sheet–rich conformers, promote rapid nucleation, and yield shorter fibrils with altered morphology. These aggregation changes are associated with enhanced β-cell stress and reduced viability in lipid-loaded amylin-expressing cells. Importantly, live-dead analysis demonstrated significantly increased cytotoxicity under combined lipid-loading and amylin-overexpression conditions, supporting a functional interaction between lipid stress and amyloidogenic burden. Conversely, the higher number of lipid droplets in amylin-overexpressing cells suggests a coordinated response to the stress induced by amylin. It reflects altered lipid metabolism and a protective adjustment aimed at reducing lipotoxicity and preventing membrane damage. This interplay between lipid droplet surfaces, amyloid structure, and cellular stress responses may have broad implications for type 2 diabetes mellitus (T2DM), a metabolic disorder where lipid droplet dynamics are altered.

Beyond improvements in glycaemic control and insulin sensitivity, fasting has been shown to promote a reproducible increase in intracellular lipid droplets across metabolically active tissues, a phenomenon previously linked to metabolic flexibility and cellular stress buffering ^74,75^. While fasting-based dietary interventions elicit systemic and cellular adaptations that may counteract key pathogenic features of T2DM, including the proteotoxic effects of amylin aggregation. It should not supplant evidence-based pharmacotherapy; these findings highlight its potential as a structured, adjunctive strategy to improve metabolic homeostasis and proteostatic capacity in selected individuals with T2DM. Continued mechanistic and longitudinal clinical research is required to delineate the long-term efficacy, safety, and regulatory pathways through which fasting modulates amylin dynamics and β-cell health.

## Materials and Methods

### Recombinant hIAPP Preparation

Recombinant human amylin (hIAPP) was produced in E. coli as a GB1-fusion construct, as previously reported in our laboratory 20. Cells were grown in 2×YT medium supplemented with ampicillin, induced at mid-log phase with 1 mM IPTG for 5 hours at 37 °C, and then harvested by centrifugation.

For purification, pellets were solubilized in 7 M guanidine hydrochloride buffer and lysed by sonication. Cleared lysates were loaded onto a Ni–NTA column under denaturing conditions, followed by native washes and on-column refolding. The GB1–hIAPP fusion protein was eluted under native conditions and dialyzed to remove imidazole.

Fusion-tag removal was done by cyanogen bromide cleavage in 0.2 N HCl for 16–20 h in the dark. The resulting hIAPP peptide was isolated by centrifugation and extensive buffer washes. The monomeric peptide was generated by dissolving the purified material in 100% HFIP (Sigma-Aldrich, CAS: 920-66-1) and removing pre-aggregated species by centrifugation. The concentration was determined in HFIP, and monomeric hIAPP was diluted into phosphate buffer immediately before use.

### Preparation of Artificial Lipid Droplets (ALD)

Pancreatic mimicking Artificial lipid droplets (ALDs) were prepared using a modified emulsification–freeze–thaw protocol previously described 21,24. For each 1 mL ALD preparation, 70 μL glycerol trioleate (Sigma-Aldrich) and 0.5 μmol (DOPC: DOPE::7:3) (Avanti Polar Lipids, Cat. No. [A80375, A80725]) were combined in a glass vial. The desired phospholipid species (25 nmol) were then added to the mixture. The lipid mixture was flash-frozen in liquid nitrogen for approximately 30 seconds, then placed under vacuum for about 4 hours to remove residual organic solvent. After drying, the lipid film was rehydrated with 950 μL PB buffer (pH adjusted to 7.4). The suspension was vortexed vigorously for at least 10 minutes to form a crude emulsion. The resulting whitish emulsion was transferred to an ultra-low-temperature freezer (−80°C) and stored until it was completely frozen. Afterward, the emulsion was rapidly thawed in a 55°C water bath, followed by 5 freeze–thaw cycles with intermittent vortexing between cycles to promote uniform droplet formation and stability.

### Isolation of Lipid Droplets from Rat

Lipid droplets were isolated from rat liver using a previously described method ^22,23,24^, with slight modifications. Briefly, Male Sprague-Dawley rats, aged 16 weeks, were anesthetized with sodium thiopental (50 mg/kg, i.p.). The abdominal cavity was opened, and the liver was perfused through the hepatic portal vein with 50 ml of ice-cold 1× PBS. The excised liver was washed three times with 1X PBS, weighed, and immediately flash-frozen in liquid nitrogen for later use. For lipid droplet isolation, the frozen liver was thawed on ice, finely chopped, and homogenized in a glass Dounce homogenizer using 1.5 volumes of 0.9 M MEPS buffer (where *M* represents the sucrose molarity; 35 mM Pipes, 5 mM EGTA, 5 mM MgSO₄, and 0.9 M sucrose, pH 7.4) containing 2× complete protease inhibitor cocktail, 4 mM PMSF, 2 µg/ml pepstatin A, and 4 mM DTT. The homogenate was centrifuged at 1,800 × *g* for 10 min at 4°C to obtain the post-nuclear supernatant (PNS). The PNS was then mixed with 1.5 volumes of 2.5 M MEPS buffer, and 15 ml of this mixture was placed at the bottom of a clear Beckman SW32 rotor tube. It was overlaid sequentially with 5 ml layers of 1.4, 1.2, 0.5, and 0 M MEPS buffers and centrifuged at 120,000 × *g* for 1 hr at 4°C. The uppermost whitish layer containing lipid droplets was collected using a 20G1 needle, flash-frozen, and stored in liquid nitrogen.

### Surface Plasmon Resonance

SPR experiments were performed on a Biacore T200 instrument (Cytiva, USA) at 25 °C using HBS-N buffer (10 mM HEPES, 150 mM NaCl, pH 7.4) as the running buffer, supplemented with 0.005% (v/v) Tween-20 to minimize nonspecific adsorption. Artificial lipid droplets (ALDs) were prepared as described above and immobilized onto an L1 sensor chip (Cytiva) designed for lipid monolayer capture. Before immobilization, the L1 chip surface was conditioned by sequential injections of 20 mM CHAPS, 40 mM octyl β-D-glucopyranoside (OG), and running buffer (each for 1 min at 30 μL/min) to remove contaminants and prepare the lipid-binding surface. ALDs were then injected at a concentration of 0.5 mg/mL in HKM buffer (120 mM potassium acetate, 1 mM MgCl₂, 50 mM HEPES, pH 7.4) at a flow rate of 5 μL/min until the desired surface coverage (∼5,000–7,000 resonance units, RU) was achieved. Unbound droplets were removed by a brief injection of 10 mM NaOH (30 s at 30 μL/min). A reference channel without lipid immobilization was used for background subtraction. hIAPP was dissolved in running buffer at concentrations ranging from 0.1 to 10 μM and injected over the ALD-coated and reference surfaces at a flow rate of 30 μL/min for 180 s (association phase), followed by 300 s of running buffer flow (dissociation phase). Between injections, the lipid surface was regenerated using 20 mM CHAPS (flow rate of 30 μL/min for 60 s), which effectively removed bound protein without destabilizing the lipid layer. Sensorgrams were processed and double-referenced (subtraction of both reference channel and blank buffer injections) using BIAevaluation software (version 4.1, Cytiva). The association rate constant (k□), dissociation rate constant (k_d), and equilibrium dissociation constant (KD) were obtained by global fitting of the sensorgrams to a 1:1 Langmuir binding model with mass-transport corrections. All experiments were performed in triplicate to ensure reproducibility.

### 15N-1H HSQC spectroscopy

Uniformly ^15^N-labelled hIAPP was expressed in *E. coli* in minimal (M9) media using ^15^NH_4_Cl (Cambridge Isotope Laboratories, Inc., Lot No. I-22394H) as the sole nitrogen source and purified by standard chromatographic procedures. The final NMR buffer consisted of 20 mM sodium phosphate, pH 7.4, and 100 mM NaCl, and the samples were adjusted to 200 µM protein concentration. Unless otherwise stated, samples were prepared in a final volume of 500 µL in 5 mm NMR tubes, supplemented with 10% (v/v) D2_22O (CAS No. 7789-20-0) for the lock and 0.02% (w/v) sodium azide to prevent microbial growth. All NMR experiments were recorded on an Avance III 750 MHz spectrometer (Bruker BioSpin, Switzerland) equipped with a cryogenic probe at 283 K. Two-dimensional ^1^H–^15^N HSQC spectra were acquired using a standard gradient-enhanced pulse sequence (Bruker pulse program *hsqcetfpgp*), with 1024 complex points in the direct dimension (F2) and 256 increments in the indirect dimension (F1), spectral widths of ∼16 ppm in ^1^H and ∼40 ppm in ^15^N, and 8 scans, and 1.5 s recycle delay. A reference HSQC spectrum of free amylin was first acquired, after which aliquots of ALDs and LDs were added sequentially, and HSQC spectra were recorded following a short incubation period of ∼10–15 min to allow equilibration. Spectra were processed in TopSpin 4.1 (Bruker BioSpin) and analyzed using NMRFAM-SPARKY ^25^. CSPs were calculated using

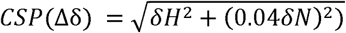

and normalized relative to the 0 µM reference spectrum to derive binding isotherms and dissociation constants (K_d_) ^26^.

### Thioflavin T (ThT) assay

Thioflavin T (ThT) dye is used to analyze the aggregation kinetics properties of amyloid fibrils. Thioflavin binds to the cross-beta sheet region of the amyloid protein and increases the fluorescence intensity in the emission region. hIAPP is to be diluted in 10mM PB Buffer (pH 7.4) to a final concentration of 100 µM. The protein is incubated at 37 °C and 120 rpm to induce aggregation. Before taking the readings, the protein sample was mixed with 100 µM ThT solution and incubated at room temperature for 10 minutes. 24-hour time-dependent Fluorescence spectra were recorded in triplicate on a Spectrofluorometer (Horiba Fluoromax-4) by exciting the samples at 450 nm and measuring the emission maxima at 482 nm ^27^. Normalized fluorescence spectra were plotted as ThT fluorescence intensity (AU) versus emission wavelength (nm). The lag time and half-time were calculated through the following equations:

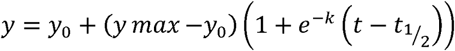

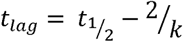

where y is the ThT fluorescence intensity at the particular point of the intensity curve, y_0_ is the ThT fluorescence intensity at t_0_ (initiation of aggregation), y_max_ is the maximum ThT fluorescence, k is the rate of aggregation, and t_1/2_ is the time to reach 50 % of y_max_ ^28^.

### Circular Dichroism (CD) spectroscopy

hIAPP was diluted to a final concentration of 50 μM in 10 mM PBS (pH 7.4). It was then incubated at 37°C and 120 rpm to induce aggregation. Time-dependent Far-UV CD spectra were recorded on a JASCO J-1500 spectropolarimeter using a 0.1 cm path-length cuvette at 25°C. CD spectra were represented as plots of ellipticity (deg/cm^2^) versus wavelength.

Following baseline correction, far-UV circular dichroism (CD) spectra were obtained under the specified experimental conditions. Yang et al.’s ^29^ reference-based technique, which links mean residue ellipticity values to fractional contributions of α-helix, β-sheet, β-turn, and random coil structures, was used to determine the secondary structure content. As previously established for protein secondary structure estimation using CD measurements, the experimental CD data were converted to mean residue ellipticity and compared with the Yang reference dataset. Using far-UV CD spectra, this method provides a quantitative, repeatable evaluation of secondary structural components.

### Atomic Force Microscopy (AFM)

The temporal changes in the morphology of hIAPP on interaction with ALDS and LDs were monitored using atomic force microscopy (AFM) (Asylum Research, Santa Barbara, CA). Ten microliters of the protein solution samples were taken and diluted to a final concentration of 100 µM. The protein solution was then spotted onto a freshly cleaved mica sheet with a smooth and uniform surface. The samples are incubated at room temperature for 20 minutes, then washed with Nuclease-Free Water and dried under vacuum for 2 hours. Imaging was performed on 2-3 randomly selected areas using a silicon nitride cantilever in tapping mode at a frequency of 300 kHz. A Frame of 5µm × 5µm was selected for analysis. Approx. 50 fibrils selected from each frame to calculate the average length and width using Igor Pro V6 software ^30^.

### Cell Culture and Generation of IAPP-GFP Stable INS1 Cell Line

For details and the CatLog number of all chemicals and kits used, kindly refer to Supplementary Table 1.

INS1 rat insulinoma cells were maintained in RPMI-1640 medium supplemented with 10% fetal bovine serum (FBS) at 37 °C in a humidified 5% CO₂ incubator. The IAPP-GFP expression cassette was cloned into a PiggyBac transposon vector using the online VectorBuilder design platform as previously reported in our laboratory. ^31^

For stable integration, cells were seeded onto gelatin-coated 24-well plates at a density of 50,000 cells/cm². After 24 hr, the culture medium was replaced with Opti-MEM (Gibco), and cells were transfected with a mixture of 300ng IAPP-GFP PiggyBac plasmid and 300 ng PiggyBac transposase helper plasmid using the Lipofectamine 3000 transfection system (Invitrogen) according to the manufacturer’s instructions. Cells were maintained in transfection medium for 14 hours, followed by replacement with complete RPMI-1640 medium for an additional 24 hours.

Transfected cells were subjected to puromycin (Sigma) selection (1 µg/mL) for 48 h. Following selection, cells were trypsinized, resuspended in FACS buffer (2.5% FBS in PBS), and sorted for GFP-positive populations using a BD FACS Aria SORP cell sorter ^32^. Sorted cells were collected into RPMI-1640 containing 20% FBS and seeded onto gelatin-coated 24-well plates. After allowing 6 h for attachment, the medium was replaced with fresh complete RPMI-1640 (20% FBS), and cells were expanded to confluency. GFP expression in the stable line was confirmed using an Olympus IX83 inverted fluorescence microscope equipped with a FITC filter.

### Calcein/Propidium Iodide (PI) Live-Dead Assay

The effect of lipid droplet–amylin interactions on the viability of IAPP-overexpressing INS1 cells was assessed using a Calcein/PI live-dead assay ^33^. Stable INS1-Amylin-GFP cells were seeded onto gelatin-coated 35-mm dishes at a density of 150,000 cells/cm². After 24 h, cells were treated with tetracycline (Sigma) (2 µg/mL) to induce Amylin-GFP expression and with oleic acid (Sigma) (100 µM) to promote lipid droplet formation. Cultures were incubated for 48 h.

Following treatment, cells were washed once and incubated in plain RPMI-1640 containing Calcein (Sigma) (5 µM) and propidium iodide (Sigma) (PI; 2 µM) for 45 min at 37 °C. Cells were then washed three times with PBS and stained with Hoechst 33342 (Sigma) (1:5000 dilution) prepared in Krebs-Ringer buffer (Himedia) for 2–3 min. Excess dye was removed by PBS washes, and cells were maintained in Krebs-Ringer buffer containing 11 mM glucose (Sigma) during imaging.

Live-cell imaging was performed on an Olympus IX83 inverted fluorescence microscope equipped with an environmental chamber. Images were acquired at 20× magnification using DAPI, FITC, and TRITC filter sets. PI-positive (dead) cells were quantified using ImageJ software and expressed as a percentage of total nuclei (Hoechst-positive cells). Graphs were generated using GraphPad Prism.

### Confocal imaging

For high-resolution imaging of IAPP localization, INS1 IAPP-GFP stable cells were seeded on 12-mm gelatin-coated coverslips and treated under the same conditions described for the Calcein/PI live-dead assay. After 48 hours of induction and oleic acid treatment, the cells were washed with PBS and fixed in 4% paraformaldehyde (PFA) (Sigma) for 15 minutes at room temperature. Fixed cells were washed and permeabilized with 0.15% Triton X-100 (Sigma) in PBS for 15 min, followed by three washes in PBS.

Cells were then blocked with 5% FBS in PBS for 2 h at room temperature. Anti-amylin primary antibody (abclonal) was added at a 1:300 dilution and incubated overnight at 4 °C. The following day, coverslips were washed with PBS and incubated with Alexa Fluor 488-conjugated goat anti-rabbit (GAR-488) secondary antibody (Invitrogen) for 4 h at room temperature. After secondary incubation, cells were washed again and stained with DAPI (1:2000 in PBS) for 10 min.

Finally, coverslips were washed thoroughly and mounted using a LipidTOX (Invitrogen) red neutral lipid stain–containing antifade mounting medium. Confocal images were acquired using a Zeiss laser-scanning confocal microscope 780 equipped with a 63× oil-immersion objective. All imaging parameters were maintained consistently across samples. All images were analyzed using ImageJ software.

### qRT–PCR Analysis

List of primers used in qRT-PCR (Table S2)

Total RNA was isolated using TRIzol® reagent (Invitrogen) according to the manufacturer’s protocol. Following phase separation with chloroform and isopropanol precipitation, RNA pellets were washed with 75% ethanol and resuspended in RNase-free water. RNA integrity and concentration were determined using a NanoDrop 2000 (Thermo Scientific). Samples with A260/280 ratios of 1.9–2.1 were used for cDNA synthesis.

cDNA was generated from 1 µg RNA using the High-Capacity cDNA Reverse Transcription Kit (Applied Biosystems). qRT–PCR was performed with PowerUp™ SYBR® Green Master Mix (Applied Biosystems) on a QuantStudio™ 6 Flex system. Reactions (20 µL) contained 0.5 µL gene-specific primers (10 µM), 2 µL cDNA (1:10 dilution), and SYBR Green Master Mix. Primers were designed using NCBI Primer-BLAST and synthesized by Eurofins Genomics.

LD Genes (Table S3) analyzed included pancreatic transcription factors (PAX6, PDX1, NKX6.1, MAFA) ^34,35,36^, lipid-metabolic regulators (PIL1, PIL2, BSC2, ATGL) ^37,38^, apoptosis markers (CASP3, CASP8, CASP9) ^39,40^, hypoxia/cytoskeleton genes (HIF1A, CYK19) ^41,42^, ER-stress markers (PERK, CHOP, ATF6, ATF4) ^43,44^, and GAPDH ^45^ as the internal control. Cycling conditions were 95 °C for 10 min, followed by 40 cycles of 95 °C for 15 s and 60 °C for 30 s. Melt-curve analysis confirmed specificity.

### Data Analysis

All qRT-PCR reactions were performed in triplicate (technical replicates) for three independent biological experiments. Gene expression was normalized to GAPDH, and relative expression levels were calculated using the 2⁻ΔΔCt method. Statistical analysis was performed using one-way ANOVA followed by Tukey’s multiple comparisons test in GraphPad Prism 9 (GraphPad Software) ^46,47^. Differences were considered significant at *p* < 0.05.

### Lipid Analysis and Data Processing

Study groups were defined based on glycated hemoglobin (HbA1c) levels according to criteria established by the World Health Organization: Healthy controls (HbA1c < 5.7%), Prediabetic individuals (HbA1c 5.7-6.4%), and Diabetic individuals (HbA1c 6.4-8.5%) ^48^.

Blood samples were collected from 30 participants: 10 healthy individuals, 10 with prediabetes, and 10 with diabetes. Serum samples were pre-aliquoted (9 µL) and stored at 4 °C immediately prior to extraction. Lipids were extracted by adding 81 µL of a 1:1 (v/v) butanol: methanol solvent containing diluted Equi-SPLASH internal standards to each aliquot. Samples were briefly vortexed and then mixed on a thermomixer at maximum speed for 15 min. Following extraction, the supernatant was transferred to LC-MS vials for subsequent analysis.

Lipidomic data were imported into Python and pre-processed by removing sample identifiers and standardizing group labels into three categories: healthy, prediabetic, and diabetic. Lipid intensities were log₁₀ transformed after adding a pseudocount of 1 to stabilize variance. Group-wise comparisons for each lipid were performed using the two-sided Mann-Whitney U test. Results were visualized using boxplots with significance annotations (*p < 0.05, **p < 0.01, ***p < 0.001; ns, not significant).

## Acknowledgments

We thank Prof. Roop Mallik and Ms. Apurwa Meghna for insightful discussions and guidance on lipid droplet–related experiments. We thank Ms. Kalyani Patil for helping with protein purification. We acknowledge the Bio-AFM, Surface Plasmon Resonance (SPR), and Confocal Laser Scanning Microscopy facilities at the Central Facility in the Department of Biosciences and Bioengineering (BSBE), Indian Institute of Technology Bombay, for access to instrumentation and technical support.

We also acknowledge the High-Field NMR (750 MHz), Circular Dichroism (CD), and Dynamic Light Scattering (DLS) facilities at IIT Bombay, funded by RIFC, IRCC, and IIT Bombay.

AKJ thanks UGC and IRCC for the fellowship. AK thanks SCAN, IIT Bombay, (DO/2023-SSAN002-003).

## References

1. P. Westermark, et al., Islet amyloid polypeptide, islet amyloid, and diabetes mellitus. Physiol. Rev. 91, 795–826 (2011).

2. G. J. S. Cooper, Amylin compared with calcitonin gene-related peptide: Structure, biology, and relevance to metabolic disease. Endocr. Rev. 15, 163–201 (1994).

3. D. L. Hay, et al., Amylin receptors: Molecular composition and pharmacology. Biochem. Soc. Trans. 32, 865–867 (2004).

4. A. Abedini, D. P. Raleigh, The role of His-18 in amyloid formation by human islet amyloid polypeptide. Biochemistry 44, 16284–16291 (2005).

5. R. Akter, et al., Islet amyloid polypeptide: Structure, function, and pathophysiology. J. Diabetes Res. 2016, 2798269 (2016).

6. G. P. Gorbenko, et al., The role of lipid–protein interactions in amyloid-type protein fibril formation. Chem. Phys. Lipids 141, 72–82 (2006).

7. F. U. Hartl, Protein misfolding diseases. Nature 475, 324–332 (2011).

8. J. F. Rivera et al., Autophagy defends pancreatic β-cells from human islet amyloid polypeptide toxicity. Cell Death Differ. 21, 343–354 (2014).

9. C. Greenhill, Protein misfolding in β-cell failure. Nat. Rev. Endocrinol. 21, 591 (2025).

10. P. G. Zavarzadeh, et al., Exploring proinsulin proteostasis: Insights into beta cell health and diabetes. Front. Mol. Biosci. 12, 1554717 (2025).

11. B. O. W. Elenbaas et al., Membrane-catalyzed aggregation of islet amyloid polypeptide is dominated by secondary nucleation. Biochemistry 61, 1465–1472 (2022).

12. J. D. Knight, et al., Phospholipid catalysis of diabetic amyloid assembly. J. Mol. Biol. 341, 1175–1187 (2004).

13. R. A. Ritzel et al., Human islet amyloid polypeptide oligomers disrupt cell coupling, induce apoptosis, and impair insulin secretion in isolated human islets. Diabetes 56, 65–71 (2007).

14. T. C. Walther, R. V. Farese, Jr., Lipid droplets and cellular lipid metabolism. Annu. Rev. Biochem. 81, 687–714 (2012).

15. M. A. Roberts, et al., Protein quality control and lipid droplet metabolism. Annu. Rev. Cell Dev. Biol. 36, 115–139 (2020).

16. C. Long, et al., Lipid droplet dynamics in type 2 diabetes and its complications: Pathophysiological insights and therapeutic options. Lipids Health Dis. 24, 284 (2025).

17. L. L. Listenberger, et al., Lipotoxicity and LD biogenesis in metabolic disease. Proc. Natl. Acad. Sci. U.S.A. 100, 3077–3082 (2003).

18. X. Tong, et al., Lipid droplets’ role in the regulation of β-cell function and β-cell demise in type 2 diabetes. Endocrinology 163, bqac007 (2022).

19. J. A. Olzmann, P. Carvalho, Dynamics and functions of lipid droplets. Nat. Rev. Mol. Cell Biol. 20, 137–155 (2019).

20. R. Dubey, et al., Recombinant human islet amyloid polypeptide forms shorter fibrils and mediates β-cell apoptosis via oxidative stress. Biochem. J. 474, 3915–3934 (2017).

21. A. R. Thiam, et al., COPI buds 60-nm lipid droplets from reconstituted water–phospholipid–triacylglyceride interfaces, suggesting a tension clamp function. Proc. Natl. Acad. Sci. U.S.A. 110, 13244–13249 (2013).

22. P. Barak, A. Rai, A. K. Dubey, P. Rai, R. Mallik, Reconstitution of microtubule-dependent organelle transport. Methods Enzymol. 540, 231–248 (2014).

23. K. Sadh, P. Rai, R. Mallik, Feeding–fasting dependent recruitment of membrane microdomain proteins to lipid droplets purified from the liver. PLoS One 12, e0183022 (2017).

24. M. Kumar, et al., Insulin activates the intracellular transport of lipid droplets to release triglycerides from the liver. J. Cell Biol. 218, 3697–3713 (2019).

25. W. Lee, M. Tonelli, J. L. Markley, NMRFAM-SPARKY: Enhanced software for biomolecular NMR spectroscopy. Bioinformatics 31, 1325–1327 (2015).

26. M. P. Williamson, Using chemical shift perturbation to characterise ligand binding. Prog. Nucl. Magn. Reson. Spectrosc. 73, 1–16 (2013).

27. H. Willander, et al., BRICHOS domains efficiently delay fibrillation of amyloid β-peptide. J. Biol. Chem. 287, 31608–31617 (2012).

28. R. Panigrahi, et al., SUMO1 hinders α-synuclein fibrillation by inducing structural compaction. Protein Sci. 32, e4632 (2023).

29. J. T. Yang, C.-S. C. Wu, H. M. Martinez, Calculation of protein conformation from circular dichroism. Methods Enzymol. 130, 208–269 (1986).

30. M. Anderson, et al., Polymorphism and ultrastructural organization of prion protein amyloid fibrils: An insight from high resolution atomic force microscopy. J. Mol. Biol. 358, 580–596 (2006).

31. M. A. Hazari, et al., Faster amylin aggregation on fibrillar collagen I hastens diabetic progression through β-cell death and loss of function. J. Am. Chem. Soc. 147, 15985–16006 (2025).

32. N. Nitta, et al., Intelligent image-activated cell sorting. Cell 175, 266–276.e13 (2018).

33. A. Kummrow, et al., Quantitative assessment of cell viability based on flow cytometry and microscopy. Cytometry A 83, 197–204 (2013).

34. M. Sander, et al., Genetic analysis reveals that PAX6 is required for normal transcription of pancreatic hormone genes and islet development. Genes Dev. 11, 1662–1673 (1997).

35. I. Artner, et al., MafA and MafB regulate genes critical to beta-cells in a unique temporal manner. Diabetes 59, 2530–2539 (2010).

36. A. E. Schaffer, et al., Nkx6.1 controls a gene regulatory network required for establishing and maintaining pancreatic beta cell identity. PLoS Genet. 9, e1003274 (2013).

37. R. Zimmermann, et al., Fat mobilization in adipose tissue is promoted by adipose triglyceride lipase. Science 306, 1383–1386 (2004).

38. D. L. Brasaemle, The perilipin family of structural lipid droplet proteins: Stabilization of lipid droplets and control of lipolysis. J. Lipid Res. 48, 2547–2559 (2007).

39. D. W. Nicholson, et al., Identification and inhibition of the ICE/CED-3 protease necessary for mammalian apoptosis. Nature 376, 37–43 (1995).

40. P. Li, et al., Cytochrome c and dATP-dependent formation of Apaf-1/caspase-9 complex initiates an apoptotic protease cascade. Cell 91, 479–489 (1997).

41. G. L. Semenza, Targeting HIF-1 for cancer therapy. Nat. Rev. Cancer 3, 721–732 (2003).

42. T. D. Pollard, Mechanics of cytokinesis in eukaryotes. Curr. Opin. Cell Biol. 22, 50–56 (2010).

43. H. P. Harding, et al., Regulated translation initiation controls stress-induced gene expression in mammalian cells. Mol. Cell 6, 1099–1108 (2000).

44. D. Ron, P. Walter, Signal integration in the unfolded protein response. Nat. Rev. Mol. Cell Biol. 8, 519–529 (2007).

45. T. Suzuki, P. J. Higgins, D. R. Crawford, Control selection for RNA quantitation. BioTechniques 29, 332–337 (2000).

46. J. W. Tukey, Comparing individual means in the analysis of variance. Biometrics 5, 99–114 (1949).

47. K. J. Livak, T. D. Schmittgen, Analysis of relative gene expression data using real-time quantitative PCR and the 2−ΔΔCt method. Methods 25, 402–408 (2001).

48. World Health Organization, Use of glycated haemoglobin (HbA1c) in the diagnosis of diabetes mellitus. Diabetes Res. Clin. Pract. 93, 299–309 (2011).

49. J. Seelig, Thermodynamics of lipid–peptide interactions. Biochim. Biophys. Acta Biomembr. 1666, 40–50 (2004).

50. J. J. W. Wiltzius, et al., Atomic structure of the cross-β spine of islet amyloid polypeptide (amylin). Protein Sci. 17, 1467–1474 (2008).

51. C. Goldsbury, et al., Amyloid fibril formation from full-length and fragments of amylin. J. Struct. Biol. 130, 352–362 (2000).

52. P. V. Dludla, et al., Pancreatic β-cell dysfunction in type 2 diabetes: Implications of inflammation and oxidative stress. World J. Diabetes 14, 130–146 (2023).

53. T. M. Ryan, et al., Phospholipids enhance nucleation but not elongation of apolipoprotein C-II amyloid fibrils. J. Mol. Biol. 399, 731–740 (2010).

54. M. M. Ouberai, et al., α-Synuclein senses lipid packing defects and induces membrane remodeling. J. Biol. Chem. 288, 9153–9168 (2013).

55. K. Shintou, J. A. Killian, Interaction of an amphipathic peptide with phosphatidylcholine/phosphatidylethanolamine mixed membranes. Biochemistry 46, 8345–8354 (2007).

56. D. H. J. Lopes, et al., Mechanism of islet amyloid polypeptide fibrillation at lipid interfaces. Biophys. J. 93, 3132–3141 (2007).

57. T. Watanabe-Nakayama, B. R. Sahoo, A. Ramamoorthy, K. Ono, High-speed atomic force microscopy reveals the structural dynamics of the amyloid-β and amylin aggregation pathways. Int. J. Mol. Sci. 21, 4287 (2020).

58. J. R. Brender, et al., Membrane-mediated self-assembly of human islet amyloid polypeptide. J. Am. Chem. Soc. 130, 6424–6429 (2008).

59. P. Cao, A. Abedini, D. P. Raleigh, Aggregation of islet amyloid polypeptide: From physical chemistry to cell biology. Curr. Opin. Struct. Biol. 23, 82–89 (2013).

60. N. Kory, R. V. Farese, Jr., T. C. Walther, Targeting fat: Mechanisms of protein localization to lipid droplets. Trends Cell Biol. 26, 535–546 (2016).

61. M. A. Welte, A. P. Gould, Lipid droplet functions beyond energy storage. Biochim. Biophys. Acta Mol. Cell Biol. Lipids 1862, 1260–1272 (2017).

62. W. Fei, et al., Molecular characterization of seipin and its mutants: Implications for seipin in triacylglycerol synthesis. J. Lipid Res. 52, 2136–2147 (2011).

63. D. Xu, et al., Rab18 promotes lipid droplet growth by tethering the ER to LDs through SNARE and NRZ interactions. J. Cell Biol. 217, 975–995 (2018).

64. M. Bosch, et al., Mammalian lipid droplets are innate immune hubs integrating cell metabolism and host defense. Science 370, eaay8085 (2020).

65. M. Bensellam, D. R. Laybutt, J.-C. Jonas, The molecular mechanisms of pancreatic β-cell glucolipotoxicity: Recent advances and future perspectives. Mol. Cell. Endocrinol. 364, 1–27 (2012).

66. C.-J. Huang, et al., High expression rates of human islet amyloid polypeptide induce endoplasmic reticulum stress mediated β-cell apoptosis, a characteristic of humans with type 2 but not type 1 diabetes. Diabetes 56, 2016–2027 (2007).

67. V. T. Samuel, G. I. Shulman, The pathogenesis of insulin resistance: Integrating signalling pathways and substrate flux. Cell Metab. 23, 785–794 (2016).

68. M. C. Petersen, G. I. Shulman, Mechanisms of insulin action and insulin resistance. Physiol. Rev. 98, 2133–2223 (2018).

69. E. Ferrannini, et al., Early metabolic markers of the development of dysglycemia and type 2 diabetes and their physiological significance. Diabetes 62, 1730–1737 (2013).

70. C. Razquin, et al., Plasma lipidomic profiling identifies a signature of progression toward type 2 diabetes. Diabetes Care 41, 184–193 (2018).

71. L. L. Listenberger, et al., Triglyceride accumulation protects against fatty acid-induced lipotoxicity. Proc. Natl. Acad. Sci. U.S.A. 100, 3077–3082 (2003).

72. E. Jarc, et al., Lipid droplets and the management of cellular stress. Yale J. Biol. Med. 92, 435–452 (2019).

73. F. Geltinger, et al., Friend or foe: Lipid droplets as organelles for protein and lipid storage in cellular stress response, aging and disease. Molecules 25, 5053 (2020).

74. A. S. Rambold, S. Cohen, J. Lippincott-Schwartz, Fatty acid trafficking in starved cells: Regulation by lipid droplet lipolysis, autophagy, and mitochondrial fusion dynamics. Dev. Cell 32, 678–692 (2015).

75. T. B. Nguyen et al., DGAT1-dependent lipid droplet biogenesis protects mitochondrial function during starvation-induced autophagy. Dev. Cell 42, 9–21.e5 (2017).

