## Supplementary for "Lipid Droplets Direct Amyloid Assembly and β-Cell Stress through Interfacial Control of Amylin Aggregation"

**Characterization of hIAPP and ALD**

The monomeric form of hIAPP or amylin is highly aggregating in nature. Due to this highly amyloidogenic nature, it is necessary to increase the solubility and stability of this peptide. The His-6 tag facilitates protein purification using Ni-NTA affinity chromatography, and GB1 is cleaved out by CNBr (Cyanogen Bromide). The purified monomeric hIAPP protein was electrophoresed on Tris-tricine SDS-PAGE, and a monomer of hIAPP showed between 6.5 and 3.5 kDa (at 4 kDa) as shown in Figure S1a. Lyophilized hIAPP monomer dissolved in 0.1 M acetic acid (pH 7.4) was subjected to MALDI-TOF to confirm the purity of the monomer preparation. A single peak was observed in the mass spectrum (at 4037 Da), as shown in Figure S1b. This protocol yielded more than 25 mg of purified fibrils/liter of 2xYT medium.

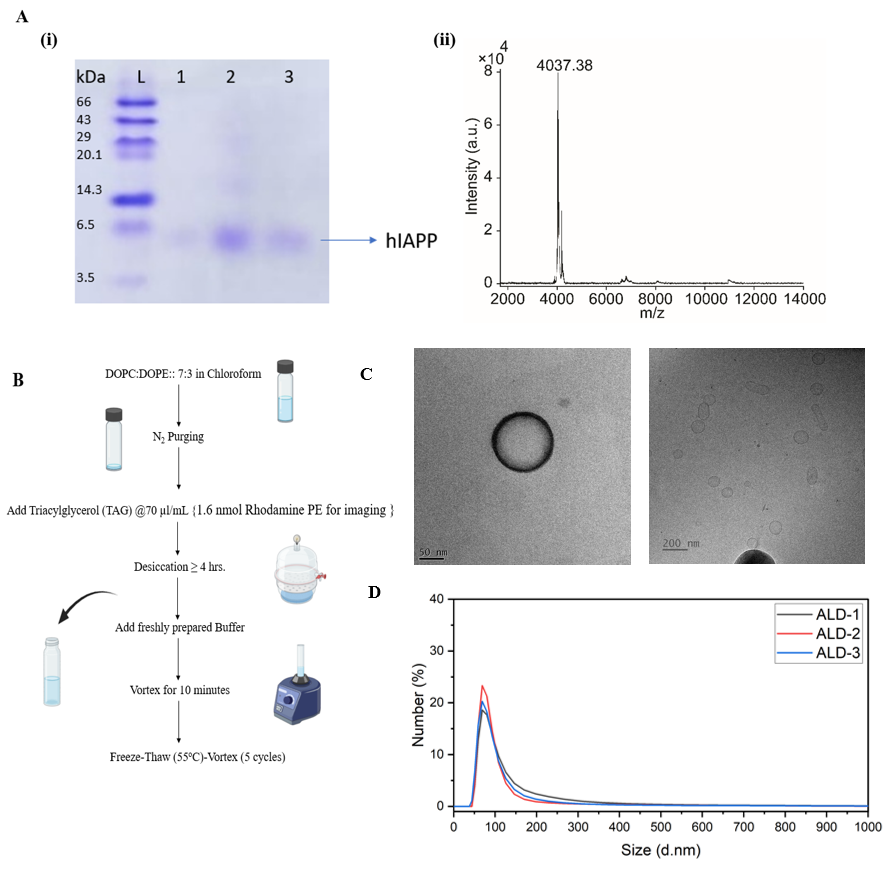

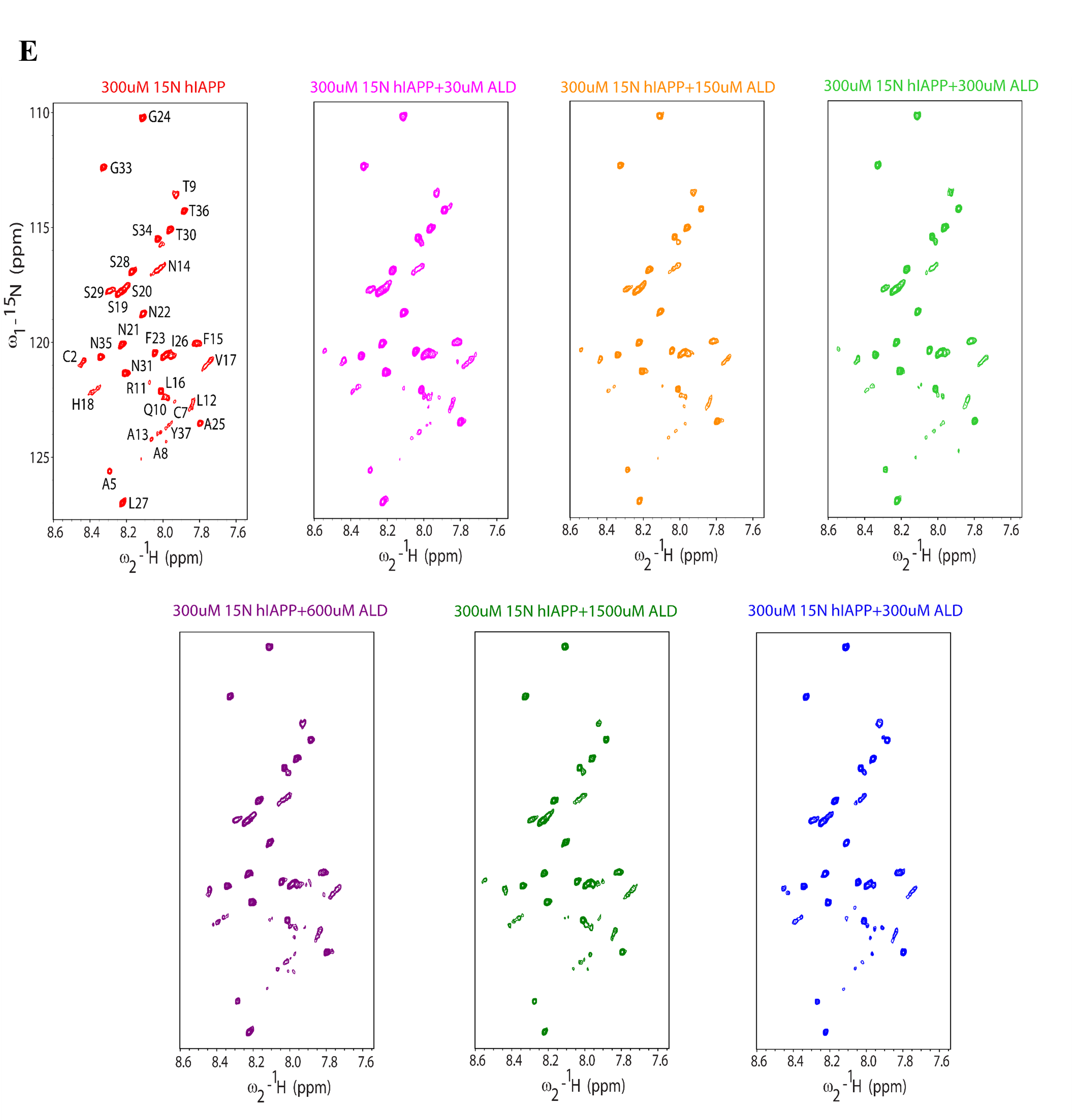

**Figure S1: Characterization of hIAPP and ALD: (A) i.** Tris-tricine SDS-PAGE of purified hIAPP. **ii.** MALDI-TOF shows pure monomeric hIAPP of 4 kDa in hexafluoroisopropanol (HFIP) solvent. **(B)** Schematic Representation showing preparation of Artificial Lipid Droplets. **(C) & (D)** Cryo-TEM image and DLS spectra showing the morphology and hydrodynamic radii of pancreatic mimicking lipid droplets (ALD), respectively. (**E)** ^15^N HSQC spectra illustrating the interaction between human IAPP (hIAPP) in Red and various concentrations of ALDs in their respective colors.

**Differential Interaction Behavior of hIAPP with Artificial and Natural Lipid Droplets**

**ALDs and LD normalization**

ALD and LD samples were normalized to OD_600_.

**LD proteolysis**

Purified lipid droplets (100 µL; OD₆₀₀ = 1) were incubated with Proteinase K (200 µg/mL final concentration) at 37 °C for 120 min. The reaction mixtures were gently mixed every 10 minutes to ensure uniform digestion. Following incubation, LDs were recovered by centrifugation at 20,000 × g for 10 minutes and subsequently resuspended in PB buffer to a final volume of 100 µL.

**Protein isolation from Lipid droplets**

Digested and untreated LD aliquots were mixed with SDS to a final concentration of 0.25% for protein extraction. Samples were heated at 60 °C and subjected to intermittent sonication (2 sec on, 5 sec off cycles for a total of 1 minute) every 20–30 min for 1 hour. Following extraction, samples were centrifuged at 20,000 × g for 10 minutes at 4 °C to isolate the protein-containing fraction.

**TLC for TG normalization**

Lipid droplets were either isolated from rat liver or reconstituted in vitro using triacylglycerol (TG) and phospholipids (DOPC: DOPE), as described in previous sections. Lipids were extracted following the standard Bligh and Dyer method (1959). Briefly, 5 µl of ALDs/LDs with an OD₆₀₀ of approximately 1 (diluted 1:500 in Milli-Q water) was mixed with 500 µl of water, followed by the addition of 2 ml methanol and 1 ml chloroform. The mixture was vortexed briefly and incubated overnight at 4°C for lipid extraction. The next day, 1 mL each of chloroform and water was added, thoroughly vortexed, and allowed to separate under gravity. The organic (lower) phase was carefully collected into a clean glass tube, and the solvent was evaporated under a gentle stream of nitrogen. Silica TLC plates were pre-cleaned with chloroform, air-dried, and heated at 100°C for 15 minutes before use. The dried lipid residues were dissolved in 100 µL of chloroform, and 10 µL of this solution was spotted onto the plate using glass capillaries. Lipid species were separated using a two-step solvent system: first, n-hexane: diethyl ether: acetic acid (60:40:4) was run to half the plate length, air-dried, and then developed fully with n-hexane: diethyl ether (59:1). After air drying, the lipids were visualized by spraying with 10% CuSO₄ in 8% H₃PO₄ and heating until bands became visible.

**Silver Staining of SDS–PAGE Gel**

Protein samples were mixed with 2× SDS–PAGE loading dye and denatured at 90 °C for 10 minutes using a dry bath. Denatured proteins were separated on a 12% SDS-polyacrylamide gel at 70 V for 3 hours. Following electrophoresis, the gel was fixed **overnight at room temperature** in a **fixation solution containing 20% (v/v) ethanol, 5% (v/v) acetic acid, and 70% (v/v) Milli-Q** with gentle shaking. The gel was then washed twice with 30% (v/v) ethanol, followed by Milli-Q for 20 minutes each.

Subsequently, the gel was incubated in a sensitization solution (0.02% sodium thiosulphate) for 15 minutes, rinsed briefly with Milli-Q (3 × 20 seconds), and then stained with 0.1% (w/v) silver nitrate for 30 minutes at 4 °C in the dark. Excess stain was removed by washing the gel twice with Milli-Q (30 seconds each). Protein bands were visualized by developing the gel in a solution containing 3% (w/v) anhydrous sodium bicarbonate and 0.05% (v/v) formaldehyde until sufficient staining intensity was achieved. The reaction was terminated by immersing the gel in 5% (v/v) glacial acetic acid.

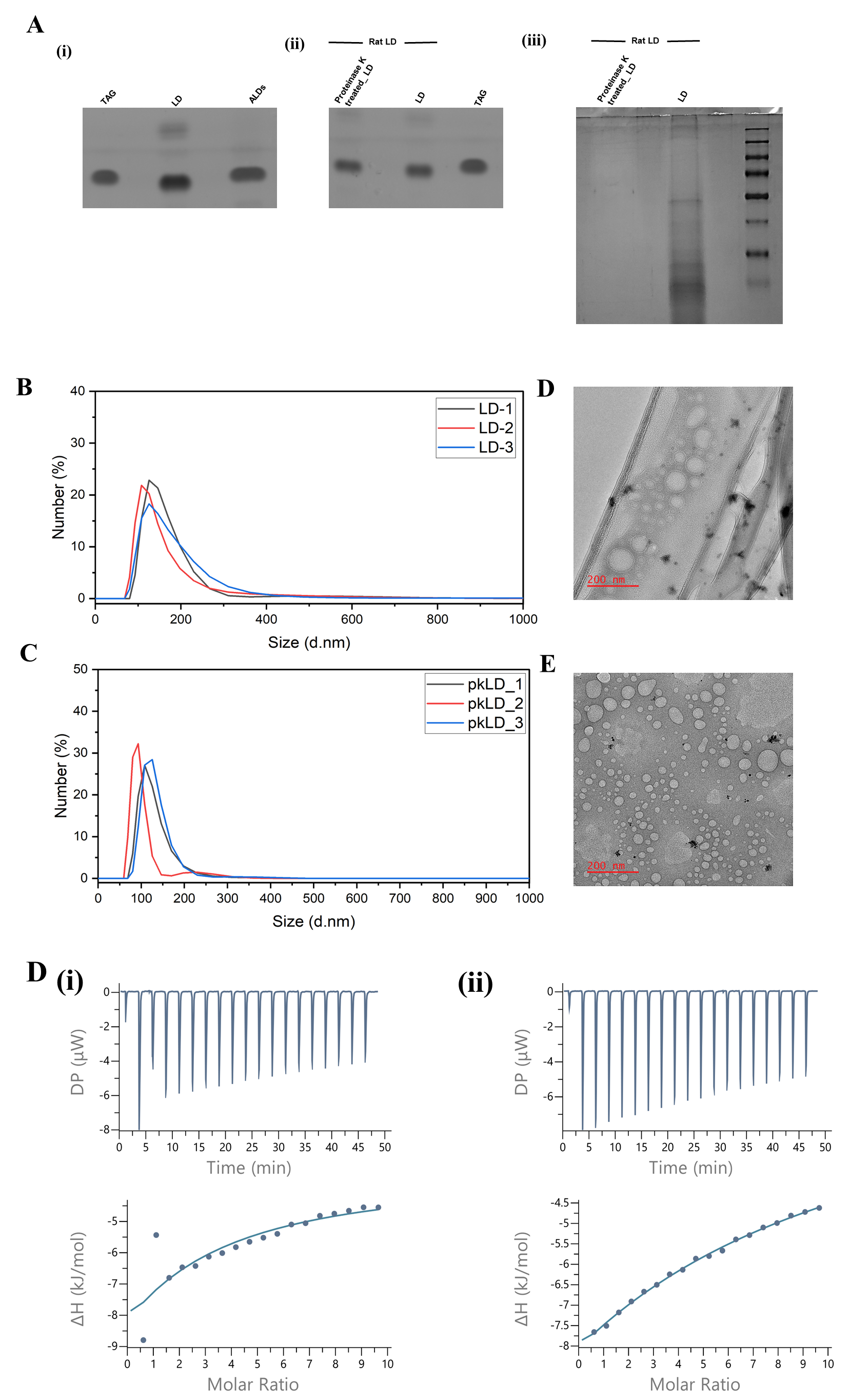

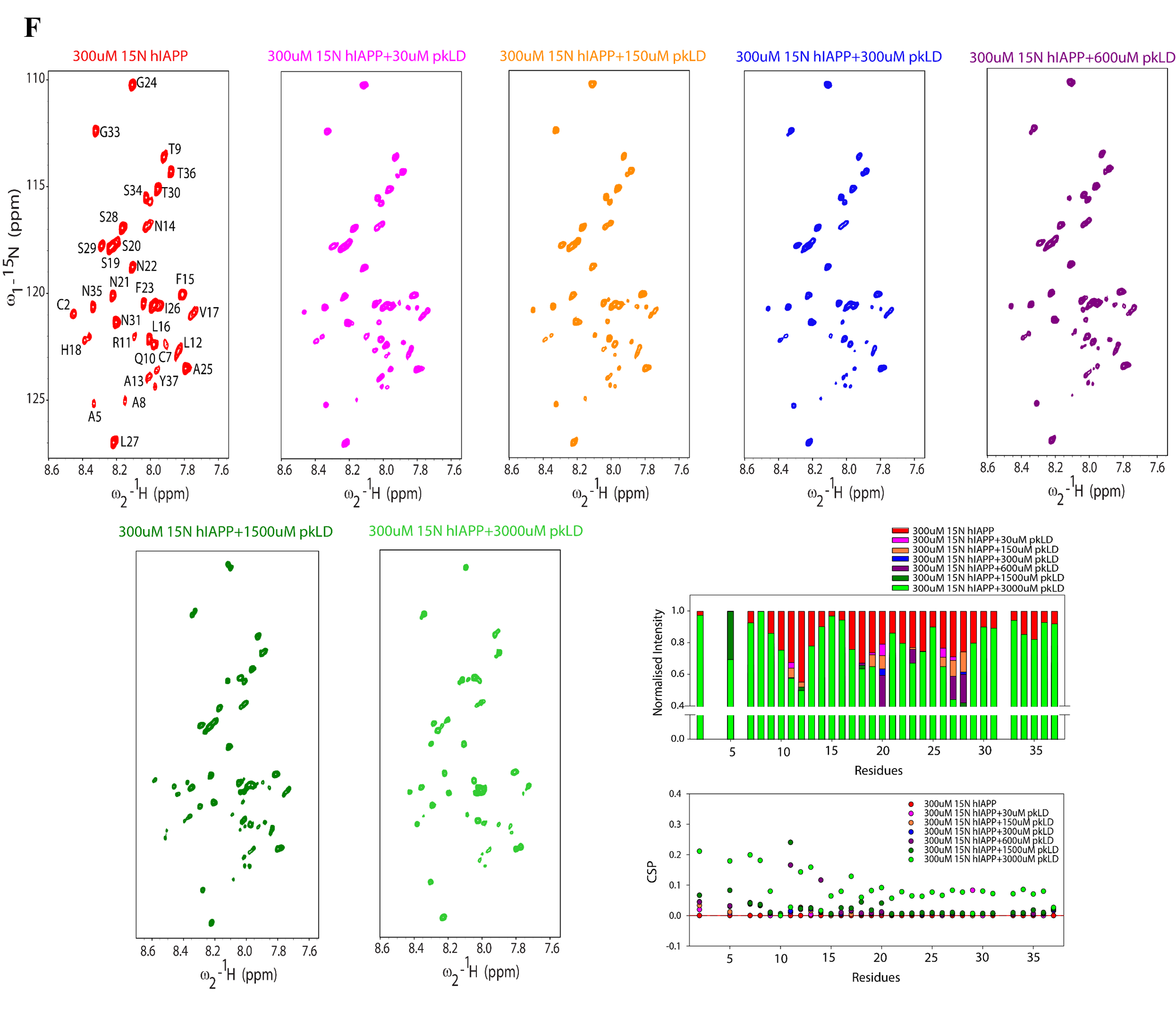

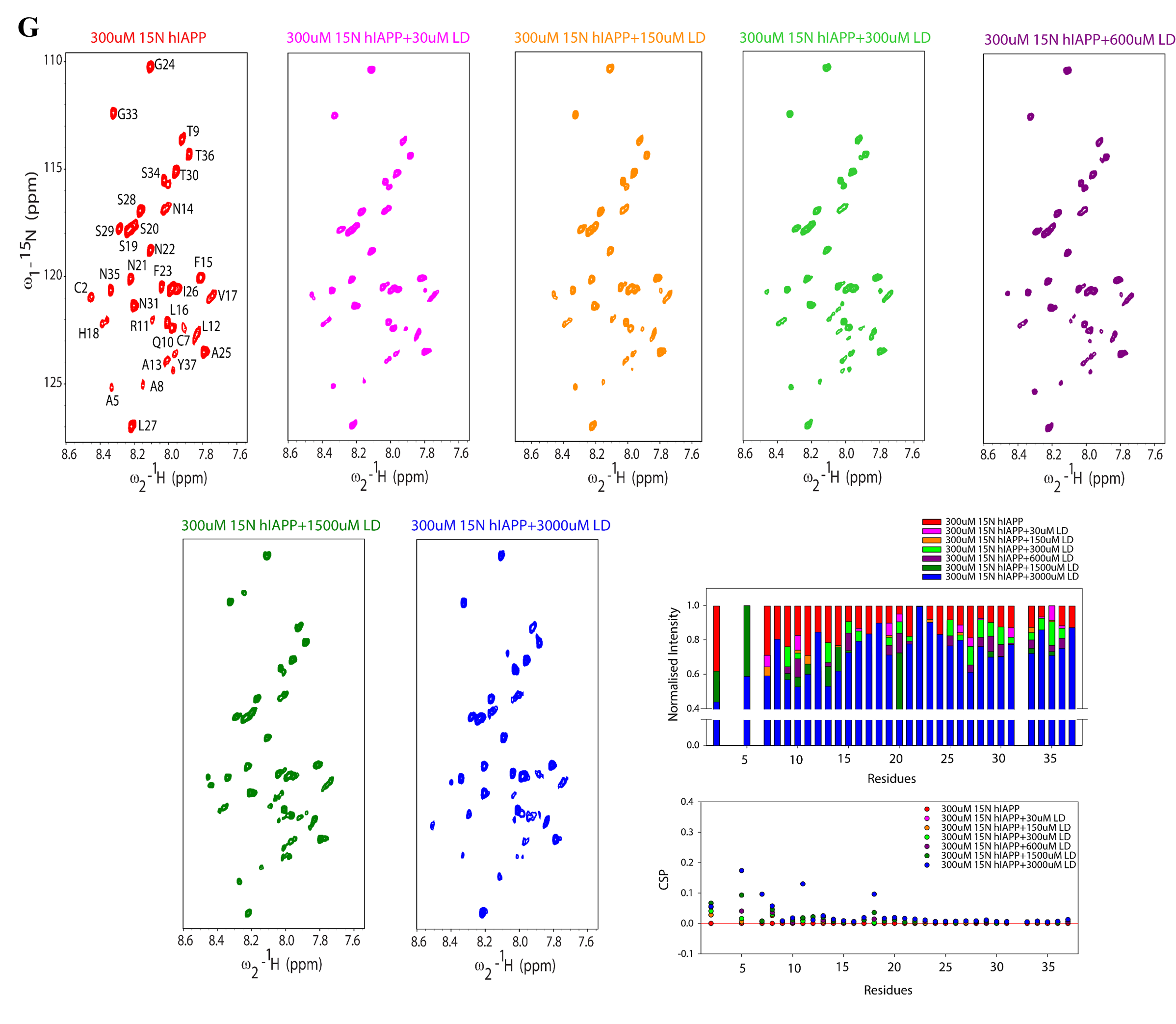

**Figure S2: (A) TLC and Silver staining for ALDs and LDs: i.** TLC showing TAG normalization for rat liver lipid droplets and artificial lipid droplets. **ii.** TLC showing TAG normalization for LD and proteinase K-treated LD. **iii.** Silver staining of protein components of purified LD before and after Proteinase K treatment. **(B), (C)** DLS and **(D), (E)** Cryo-TEM image showing hydrodynamic radii and morphology of LD and pkLD, respectively. **(F) & (G)** ^15^N HSQC spectra illustrating the interaction between human IAPP (hIAPP) in Red and various concentrations of pkLD and LD in their respective colors. Bar and dot plot of Normalized Intensity and CSP, respectively, in the lower panel.

**Live-Dead Assay of Amylin-GFP Stable INS1 Cell Line**

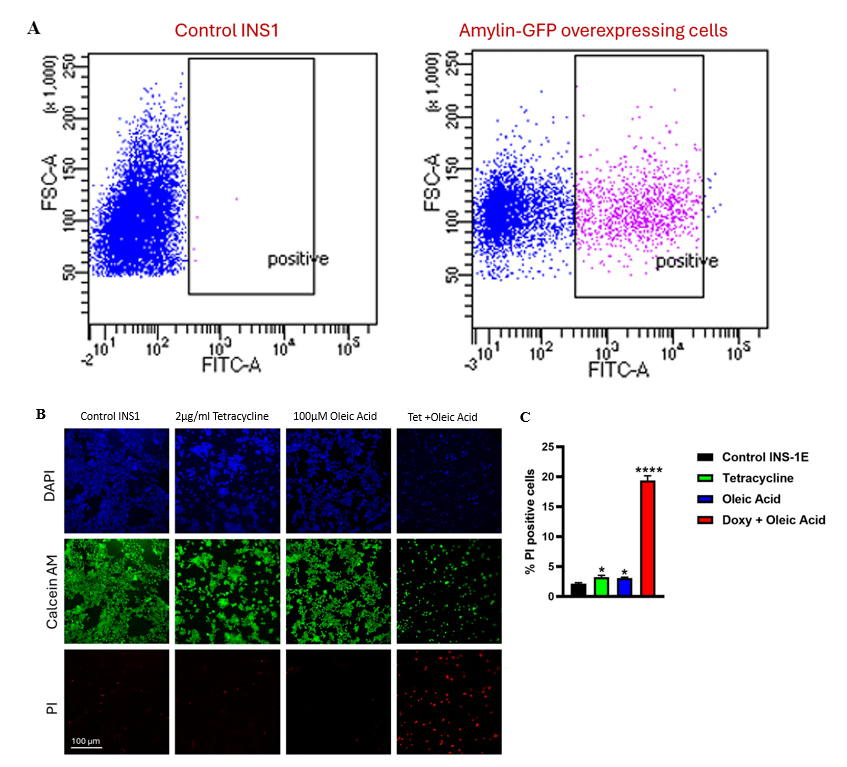

**Figure S3: (A) Flow cytometric analysis of GFP expression in INS-1E cells:**
Representative flow cytometry dot plot showing forward scatter area (FSC-A; y-axis), reflecting relative cell size, versus FITC-A fluorescence intensity (x-axis, logarithmic scale). Blue dots represent control INS-1E cells lacking GFP expression, while pink dots represent INS-1E cells overexpressing GFP. The rectangular gate indicates the GFP-positive population, defined based on the fluorescence distribution of control cells.

**(B) Representative fluorescence microscopy images showing nuclear staining, GFP expression, and cell death in INS-1 cells under different treatment conditions:** The top row shows nuclear staining (blue; DAPI), the middle row shows GFP fluorescence (green), and the bottom row shows propidium iodide (PI) staining (red), indicating membrane-compromised/dead cells. Control and single-treatment conditions display preserved nuclear integrity, with robust GFP signal and minimal PI positivity, whereas the combined-treatment condition shows a marked reduction in cell density and a pronounced increase in PI-positive cells. Scale bar: 100 µm.

**(C) Live Dead Assay:** Bar graph showing the percentage of propidium iodide (PI)–positive INS-1 cells under control conditions, treatment with tetracycline (2 µg/ml), oleic acid (100 µM), or their combination. Control cells display low basal PI positivity, while doxycycline or oleic acid alone induces only a modest increase in cell death. In contrast, combined treatment with doxycycline and oleic acid results in a significant increase in PI-positive cells. Data are presented as mean ± SEM from independent experiments. *p<0.05

**Supplementary Table 1**

**Reagents and Kits Used in the Study**

| **Reagent type or resource** | **Description** | **Source** | **Identifiers** |
| --- | --- | --- | --- |
| Cell line | INS1E | Gifted by Prof. Claes Wollheim and Prof. Pierre Maechler from the University Medical Centre, Geneva, Switzerland |  |
| Plasmids | PiggyBac transposon vector system | Synthesized using vector builder tool |  |
| Transfection system | Lipofectamine 3000 | Invitrogen | L3000001 |
| Transfection media | Opti-MEM | Gibco | 31985062 |
| RPMI-1640 | cell culture media (CCM) | Gibco | 31800-022 |
| no glucose RPMI-1640 | CCM | HiMedia | AT150 |
| fetal bovine serum (FBS) | CCM supplement | Gibco | 10270-106 |
| HEPES | CCM supplement | Gibco | 15630080 |
| sodium pyruvate | CCM supplement | Gibco | 11360039 |
| 2-mercaptoethanol | CCM supplement | Gibco | 21985023 |
| DPBS | phosphate buffer saline | Gibco | 21600010 |
| Krebs-Ringer Bicarbonate buffer | KRBH buffer | Himedia | TL1143 |
| Calcein AM | dye | Invitrogen | C1430 |
| propidium iodide (PI) | dye | Sigma | P4170 |
| Tetracycline | Drug | Sigma | T3258 |
| Puromycin | Drug | Sigma | P8833 |
| Oleic acid | Drug | Sigma | O1383 |
| Paraformaldehyde | Reagent | Sigma | 158127 |
| Triton X-100 | Reagent | Sigma | T8787 |
| Anti-amylin | Primary antibody | Abclonal | A24014 (1:300) |
| Hoechst 33342 | Dye | Sigma | AMBH9A260690 |
| LipidTOX red neutral lipid stain | Dye | Invitrogen | H34476 |
| Alexa Fluor 488–conjugated goat anti-rabbit (GAR-488) | Dye | Invitrogen | A-11008 |

**Supplementary Table 2**

**List of primers used in qRT-PCR:**

| **S. No.** | **Primer** | **Sequence** |
| --- | --- | --- |
| 1. | CHOP | Forward: GAGAGAGGCGATGTTCCAGA  Backward: CCTCTTCGTTTCCTGGGGAT |
| 2. | PERK | Forward: TGAGGAGAAGAGGGGAGTGT  Backward: AGAAAGGGCTGCTTCGTACT |
| 3. | IRE1 | Forward: AGAGCCAGCACAGTAACACT Backward: ATTGGGGTCTGGGAGGAAAG |
| 4. | ATF6 | Forward: AGCCCCTCATTAACACGACA Backward: AGAATTCGAGCCCTGTTCCA |
| 5. | Caspase 3 | Forward: AAAGCACTGGAATGACATC  Backward: CGCATCAATTCCACAATTTC |
| 6. | Caspase 8 | Forward: CTACAGGGTCATGCTCTATC  Backward: ATTTGGAGATTTCCTCTTGC |
| 7. | Caspase 9 | Forward: CTCTACTTTCCCAGGTTTTG  Backward: TTTCACCGAAACAGCATTAG |
| 8. | MAFA | Forward: CTTCAGCAAGGAGGAGGTCA  Backward: CCCGCCAACTTCTCGTATTT |
| 9. | NKX6.1 | Forward: GCCAGCAGATCTTCGCCCTG  Backward: GCTGCCTCCGCTGGATTTGT |
| 10. | PAX6 | Forward: AGGCACGGTATCAGTTGGAA  Backward: CAACCACATGAGCCAACACA |

**Supplementary Table 3**

**List of Primers associated with Lipid-Droplets present in Pancreatic β-Cells**

| **S. No.** | **Primer** | **Sequence** |
| --- | --- | --- |
| 1. | Perilipin 2 | Forward: TGCCTATTCTGAACCAGCCA  Backward: CGCCATCAGACACTTCCTTG |
| 2. | Perilipin 1 | Forward: TTGGGAAGCATCGAGAAGGT  Backward: GCTGGTGTGAGGTGTAGGAT |
| 3. | Perilipin 5 | Forward: GTGCCTACAACTCAGCCAAG  Backward: CGGTAGCCATTCTCCAGACA |
| 4. | PNPLA 2/ATGL | Forward: TCGCACTTTAGCTCCAAGGA  Backward: GTTGGTGGAGCTGTCTTGTG |
| 5. | BSCL2 | Forward: AGAAACCAGAGAAGCAGCCT  Backward: AGGGAACAAGGGAAGAGGTG |

**Material Method**

**Surface Plasmon Resonance**

SPR experiments were performed on a Biacore T200 instrument (Cytiva, USA) at 25 °C using HBS-N buffer (10 mM HEPES, 150 mM NaCl, pH 7.4) as the running buffer, supplemented with 0.005% (v/v) Tween-20 to minimize nonspecific adsorption. Artificial lipid droplets (ALDs) were prepared as described above and immobilized onto an L1 sensor chip (Cytiva) designed for lipid monolayer capture. Before immobilization, the L1 chip surface was conditioned by sequential injections of 20 mM CHAPS, 40 mM octyl β-D-glucopyranoside (OG), and running buffer (each for 1 min at 30 μL/min) to remove contaminants and prepare the lipid-binding surface. ALDs were then injected at a concentration of 0.5 mg/mL in HKM buffer (120 mM potassium acetate, 1 mM MgCl₂, 50 mM HEPES, pH 7.4) at a flow rate of 5 μL/min until the desired surface coverage (~5,000–7,000 resonance units, RU) was achieved. Unbound droplets were removed by a brief injection of 10 mM NaOH (30 s at 30 μL/min). A reference channel without lipid immobilization was used for background subtraction. hIAPP was dissolved in running buffer at concentrations ranging from 0.1 to 10 μM and injected over the ALD-coated and reference surfaces at a flow rate of 30 μL/min for 180 s (association phase), followed by 300 s of running buffer flow (dissociation phase). Between injections, the lipid surface was regenerated using 20 mM CHAPS (flow rate of 30 μL/min for 60 s), which effectively removed bound protein without destabilizing the lipid layer. Sensorgrams were processed and double-referenced (subtraction of both reference channel and blank buffer injections) using BIAevaluation software (version 4.1, Cytiva). The association rate constant (kₐ), dissociation rate constant (k_d), and equilibrium dissociation constant (KD) were obtained by global fitting of the sensorgrams to a 1:1 Langmuir binding model with mass-transport corrections. All experiments were performed in triplicate to ensure reproducibility.
